# Inverse FoldDir: Structure-conditioned Protein Sequence Design by Dirichlet Flow Matching

**DOI:** 10.64898/2026.09.06.749733

**Authors:** Alp Tartici, Mihajlo Stojkovic, Anru Tian, Michael C. Jewett, Russ B. Altman, Bruce J. Wittmann

**Author notes:** Corresponding authors: Russ B. Altman and Bruce J. Wittmann. Additional author contact: Alp Tartici. This work was initiated while Alp Tartici was an intern in the Office of the Chief Scientific Officer at Microsoft.

## Abstract

Protein engineering has important implications in the bioeconomy, enabling applications in materials, medicine, and energy. A key challenge is designing protein sequences that have a specific form and function. Protein inverse folding seeks to address this challenge by identifying amino acid sequences compatible with a desired protein backbone. This task is central to protein redesign and can provide a sequence-design capability for de novo backbones produced by structure-generation methods. Ideally, inverse folding can provide diverse sequence alternatives, fixed residues or motifs, soft biochemical preferences at selected positions, and candidates that remain experimentally useful. We developed Inverse FoldDir, a controllable inverse-folding method that performs iterative denoising on the amino acid probability simplex. Given a backbone structure, the model updates all positions jointly through a learned Dirichlet flow, supporting full sequence generation, fixed-residue inpainting, and user-defined soft residue priors. On the held-out CATH 4.2 test set, Inverse FoldDir achieved a mean TM-score of 84.5 (on a 0-100 scale) and a mean C*α* RMSD of 1.76 Å, compared with 83.3 and 1.86 Å, respectively, for ESM-IF1, the strongest evaluated baseline on both metrics. Denoising trajectory analyses showed that positions commit at different rates and that some residues change identity late in generation, illustrating whole-sequence refinement rather than one-shot prediction or irreversible sequential decoding. We experimentally tested Inverse FoldDir in an anti-GFP nanobody redesign task, where two of 35 redesigned sequences retained reproducible sfGFP-binding signal across independent assay runs with approximately 43% sequence divergence from the native nanobody. Inverse FoldDir is a structure-conditioned protein redesign method that combines structural recovery, user control, experimental validation, and a natural route toward future property-guided sampling.

## 2 Introduction

Protein engineering often starts with a backbone structure. A designer may want to preserve a scaffold, redesign a protein family member, alter an interface, stabilize a fold, or generate sequences for a newly designed backbone. Inverse folding addresses this problem by asking which amino acid sequences are compatible with a specified protein backbone [1, 2]. This task is central to scaffold design, enzyme redesign, antibody and nanobody engineering, fold stabilization, and structure-guided protein optimization [3, 4, 5]. It is also increasingly important as structure-generation methods produce novel backbones for which no natural sequence exists [6, 7].

Protein design campaigns rarely begin with either no information or a complete target sequence. Some positions may need to remain fixed, such as catalytic residues, disulfide cysteines, known interface hot spots, glycine/proline turns, or residues from a validated motif [3]. Other positions may only have a soft biochemical preference. For example, a design may require solvent-exposed positions to favor polar residues, a candidate metal-binding site to favor histidine, cysteine, aspartate, or glutamate, or a putative nucleic-acid-binding surface to favor basic or polar residues [8]. These requirements motivate combining residue-level control with sequence-wide refinement, so that designable positions can adapt to both user preferences and the evolving sequence context.

Autoregressive, one-shot, and diffusion-style inverse-folding methods have advanced structure-conditioned sequence design. Autoregressive models such as ESM IF [2], ProteinMPNN [6], and FAMPNN [9] generate residues sequentially and can produce strong designs, but the generation process is path-dependent. ProteinMPNN supports fixed residues and amino-acid biases, including position-specific preferences [6]. These biases modify the distribution when a residue is sampled and can influence subsequent choices through the sampled identity, but previously sampled residues are not revisited within the same decoding pass. One-shot models such as PiFold [10] avoid sequential path dependence but predict residue distributions in a single pass rather than iteratively revising them.

Diffusion and flow-based approaches provide an alternative through a continuous full-sequence representation. In Inverse FoldDir, fixed residues are clamped to exact identities, whereas soft preferences enter as initial amino-acid probability distributions at selected positions. Because the model conditions on the evolving sequence state at every update, a preference can influence surrounding predictions, which can in turn inform later updates at the preferred position. This permits reciprocal adaptation among designable positions without committing to residue identities early. Soft preferences guide initialization rather than enforce the final residue choice. The same representation provides a natural route toward future property-guided sampling: when differentiable scoring functions are available, generation can in principle be nudged toward desired sequence-level or structure-level properties rather than relying only on post hoc filtering. Related guidance approaches have been used in DNA promoter sequence design [11, 12].

We developed Inverse FoldDir as a structure-conditioned generative model that iteratively denoises amino-acid probability distributions across the full protein sequence. We evaluate Inverse FoldDir in four ways. First, we show how the denoising trajectory updates residue probabilities across the full sequence. Second, we benchmark structural recovery on held-out CATH proteins against established inverse-folding methods. Third, we characterize training components and sequence-level behavior. Fourth, we test selected designs in an anti-GFP nanobody binding assay.

This study makes six contributions: (1) a Dirichlet-flow-matching formulation for protein inverse folding; (2) a controllable design workflow supporting full-sequence generation, fixed-residue inpainting, and soft residue-prior conditioning; (3) an interpretable analysis of whole-sequence denoising, showing that residue positions resolve at different rates and can revise their identities during generation; (4) improved structural self-consistency on the CATH benchmark relative to established inverse-folding methods; (5) experimental validation through anti-GFP nanobody sequence redesign to demonstrate retained activity; and (6) a public software implementation with documentation and example workflows for practical protein-design use.

## 3 Results

### 3.1 Inverse FoldDir provides a controllable workflow for protein inverse folding

We trained Inverse FoldDir as a structure-conditioned generative model that combines Dirichlet flow matching with an SE(3)-equivariant residue graph encoder. Given a protein backbone, the model constructs a residue graph and iteratively denoises all amino-acid distributions on the simplices simultaneously (Figure 1A). Inverse FoldDir provides three complementary design modes. In full generation, all residues are initialized from an uninformative amino-acid Dirichlet distribution and are jointly refined into a complete sequence, making this mode suitable for designing sequences for backbones without known native sequences (Figure 1B). In fixed-residue inpainting, user-specified residues remain fixed throughout generation while all remaining positions are redesigned, enforcing explicit preservation of catalytic residues, disulfide bonds, interaction hotspots, or other experimentally validated motifs. In soft residue-prior conditioning, users initialize selected positions with custom amino-acid probability distributions rather than fixed identities, allowing residue-class preferences, such as favoring polar residues on solvent-exposed surfaces or metal-coordinating residues within candidate binding sites, without constraining the model to a single amino acid (Figure 1C).

**Figure 1:**
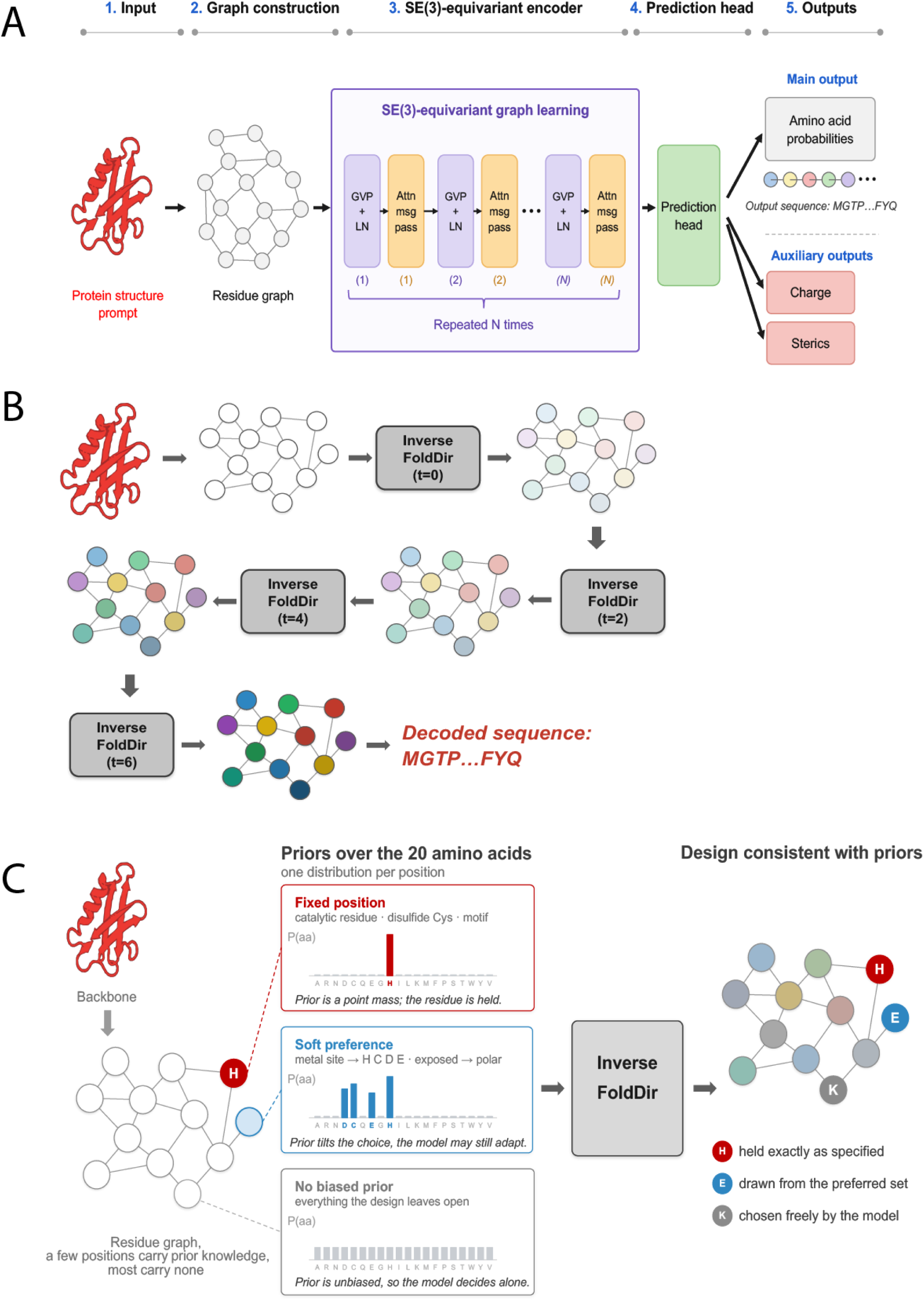
Inverse FoldDir architecture and controllable generation workflow. (A) The Inverse FoldDir model architecture consists of an SE(3)-equivariant graph representation of the backbone structure and an equivariant encoder involving interleaved Geometric Vector Perceptron layers, layer normalization, and attention-based message passing, followed by multitask predictions of denoised amino-acid probabilities and the charge and steric characteristics of each residue. The amino-acid probabilities define the Dirichlet vector field used during generation. (B) Inverse FoldDir generates sequences by iteratively updating randomly sampled probability distributions using the Dirichlet vector field. For simplicity, the illustration shows four update steps, whereas the deployed version uses 20 update steps. (C) The Inverse FoldDir framework allows users to integrate prior knowledge at desired positions by fixing residue identities and/or initializing probability distributions with soft preference biases.

Inverse FoldDir is distributed as an open-source software package with model weights, documentation, and example notebooks that demonstrate full-sequence generation, constrained redesign, candidate ranking by structural recovery, and downstream design workflows, enabling direct application by experimental and computational protein designers.

### 3.2 Iterative denoising exposes residue-level uncertainty and allows late sequence revision

Unlike existing methods, Inverse FoldDir was trained to predict denoised amino-acid probabilities at different noise levels and time points. These probabilities define the Dirichlet vector field used to update the state on the probability simplex [11] (Figure 1A). To better understand how this training paradigm constructs protein sequences, we examined the denoising trajectory for a representative bromodomain structure (PDB: 2D9E, chain A). Rather than directly predicting a final sequence, the model iteratively refines amino-acid probability distributions across all residue positions (Figure 2). This trajectory provides a window into how sequence uncertainty is resolved during inverse folding and reveals how different regions of a protein commit to their final identities over time.

**Figure 2:**
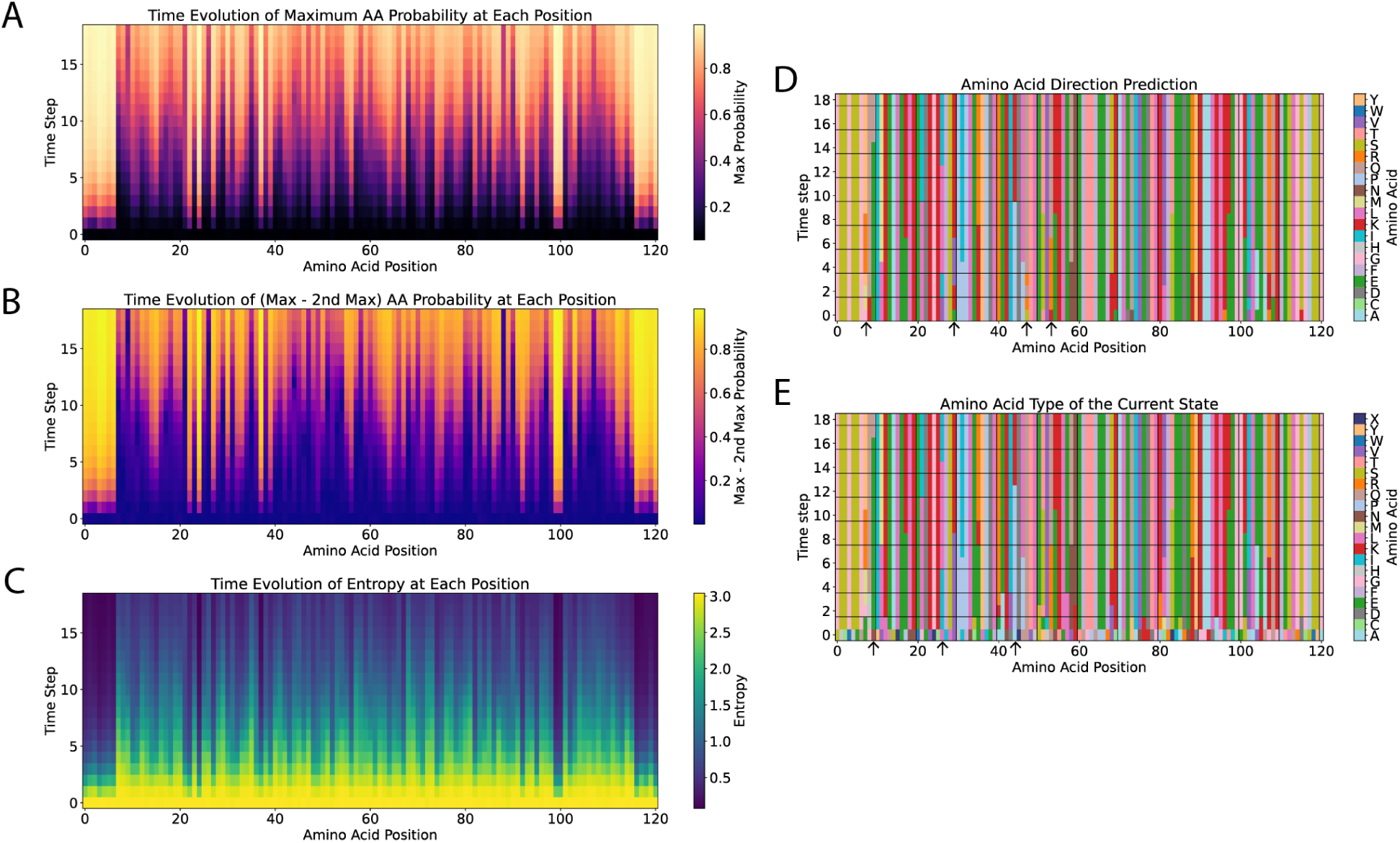
Per-position trajectory updates for a representative example, PDB structure 2D9E, chain. **A.** The x-axis denotes amino-acid position and the y-axis denotes the time step, with higher time steps indicating later stages of denoising. (A) The maximum assigned probability increases at different rates for different positions. (B) The margin between the most probable and second-most-probable amino acids varies among positions. (C) The entropy of the probability distribution decreases steadily over time, but at different rates across positions. (D) Inverse FoldDir can revise its predicted amino-acid probabilities, thereby changing the resulting Dirichlet flow direction and reversing the course of the trajectory for a position based on how neighboring positions have evolved. These changes can occur during late stages of denoising and sometimes occur up to four times. Black arrows indicate positions where the resulting flow direction changes multiple times. (E) The most likely amino acid at a given position can change during late stages of denoising, as highlighted by the positions marked with black arrows.

As denoising progresses, the maximum predicted probability for the most likely amino acid increases across all positions, indicating progressively greater confidence in the generated sequence (Figure 2A). This increase, however, was highly heterogeneous. Some residues rapidly converged toward a single amino-acid identity during the early stages of denoising, whereas others remained uncertain until much later in the trajectory, with minimal differences between the most-and second-most-probable amino acid probabilities (Figure 2B). Consistent with this observation, the entropy of the amino-acid distributions decreased throughout generation but did so at markedly different rates across residue positions (Figure 2C), suggesting that the model resolves sequence identity asynchronously rather than uniformly across the protein.

Importantly, several residue positions in this representative structure changed their most probable amino-acid identity after earlier denoising steps (Figure 2D,E). These late revisions demonstrate that residue identities remain flexible while additional sequence context develops, allowing neighboring positions to co-evolve throughout generation. These trends also generalized across the 1,120 proteins in the test set. The maximum predicted probability on the simplex generally increased (Figure 3A), while the entropy of the probability distributions generally decreased as denoising progressed (Figure 3C). The margin between the most likely and second-most-likely amino acids at the final simplex state remained highly heterogeneous (Figure 3B). The resulting Dirichlet flow direction changed during generation for more than one-quarter of the positions, and convergence did not occur until the final steps (Figure 3D). The flow direction defined by the predicted amino-acid probabilities changed more than once for an average of 7% of amino-acid positions (Figure 3E), and an average of 4% of predicted residue changes occurred during the second half of the denoising process, with a broad distribution across proteins (Figure 3F). Positions with unresolved experimental coordinates showed the greatest frequency of predicted-identity changes among the structural categories evaluated [13] (Figure S1A). More solvent-exposed positions also showed a greater tendency to update their predicted probabilities along the denoising trajectory (Figure S1B). Together, these observations show that Inverse FoldDir maintains a mutable whole-sequence representation until generation is complete rather than committing residues irreversibly at early stages. Residues therefore do not converge independently. Instead, they co-evolve as information propagates across the sequence during denoising.

**Figure 3:**
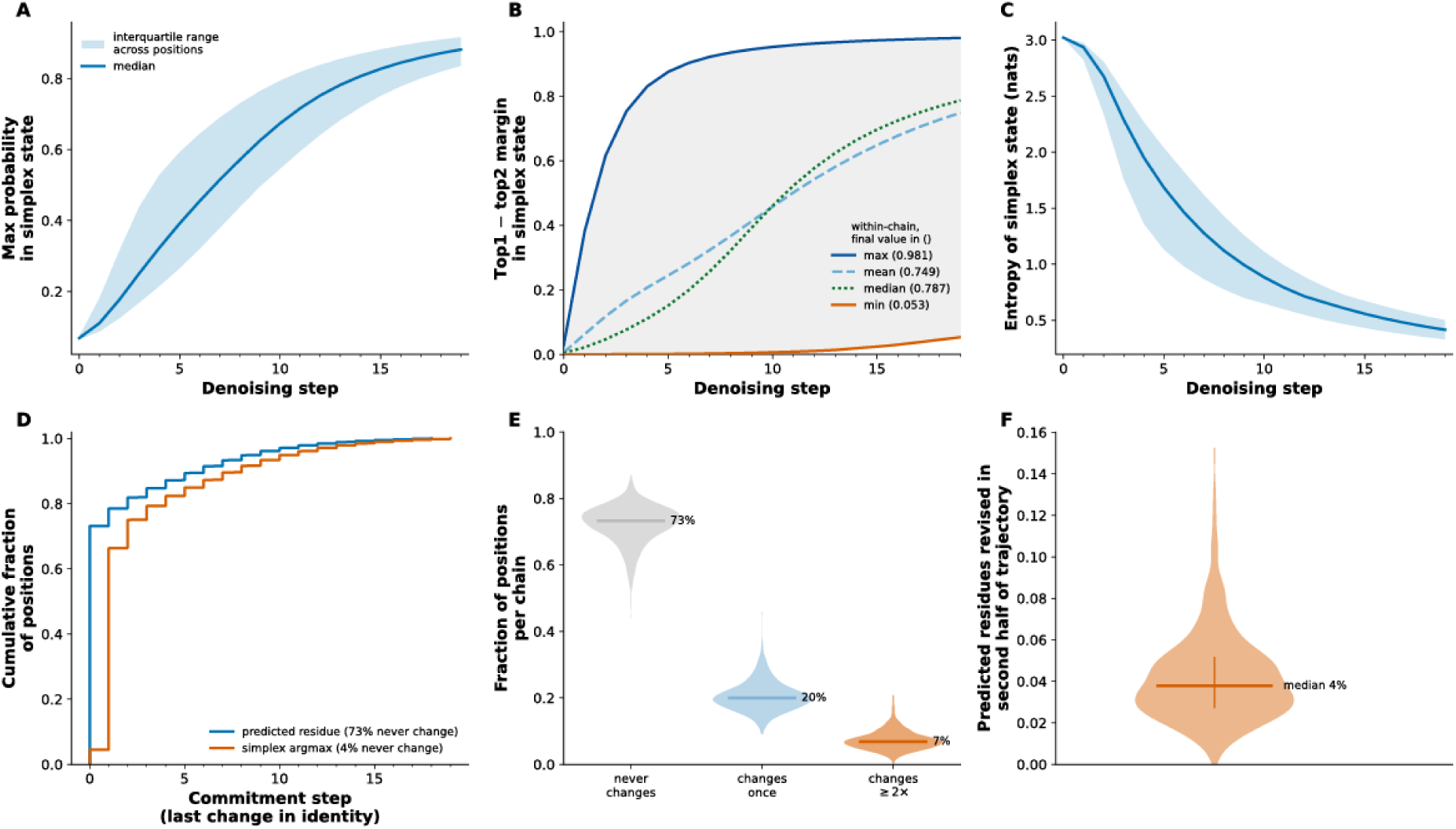
Dirichlet-flow denoising behavior across the test set. (A) The maximum probability assigned to each position increases as the probability distribution sharpens. (B) The difference between the probabilities of the most likely and second-most-likely amino acids at the final step varies across positions, demonstrating heterogeneity in prediction confidence. (C) The entropy of the simplex probability distributions decreases as the distributions sharpen. (D) On average, 27% of positions change their resulting Dirichlet flow direction during denoising (blue line), and the most likely amino acids on the simplices continue to change until late update steps (orange line). (E) Percentage of positions grouped by the number of times the flow direction changes. (F) Percentage of positions receiving updates during the second, later half of denoising.

We also examined residue-level prediction patterns in the final generated sequences. Recovery was not uniform across amino acids or physicochemical groups. Predictions frequently remained within the same physicochemical class even when the exact native residue was not recovered. Glycine and proline showed particularly high identity-level recovery (Supplementary Figure S2).

### 3.3 Inverse FoldDir improves structural recovery on held-out CATH backbones

We evaluated structural self-consistency by folding each generated sequence with ESMFold [14] and comparing the predicted structure with the input backbone using TM-score [15] and C*α* RMSD (Figure 4A). We performed the benchmark on the held-out CATH 4.2 test set [1] using a shared refolding and structure-comparison pipeline. All baseline methods were evaluated using their publicly released pretrained checkpoints without retraining.

**Figure 4:**
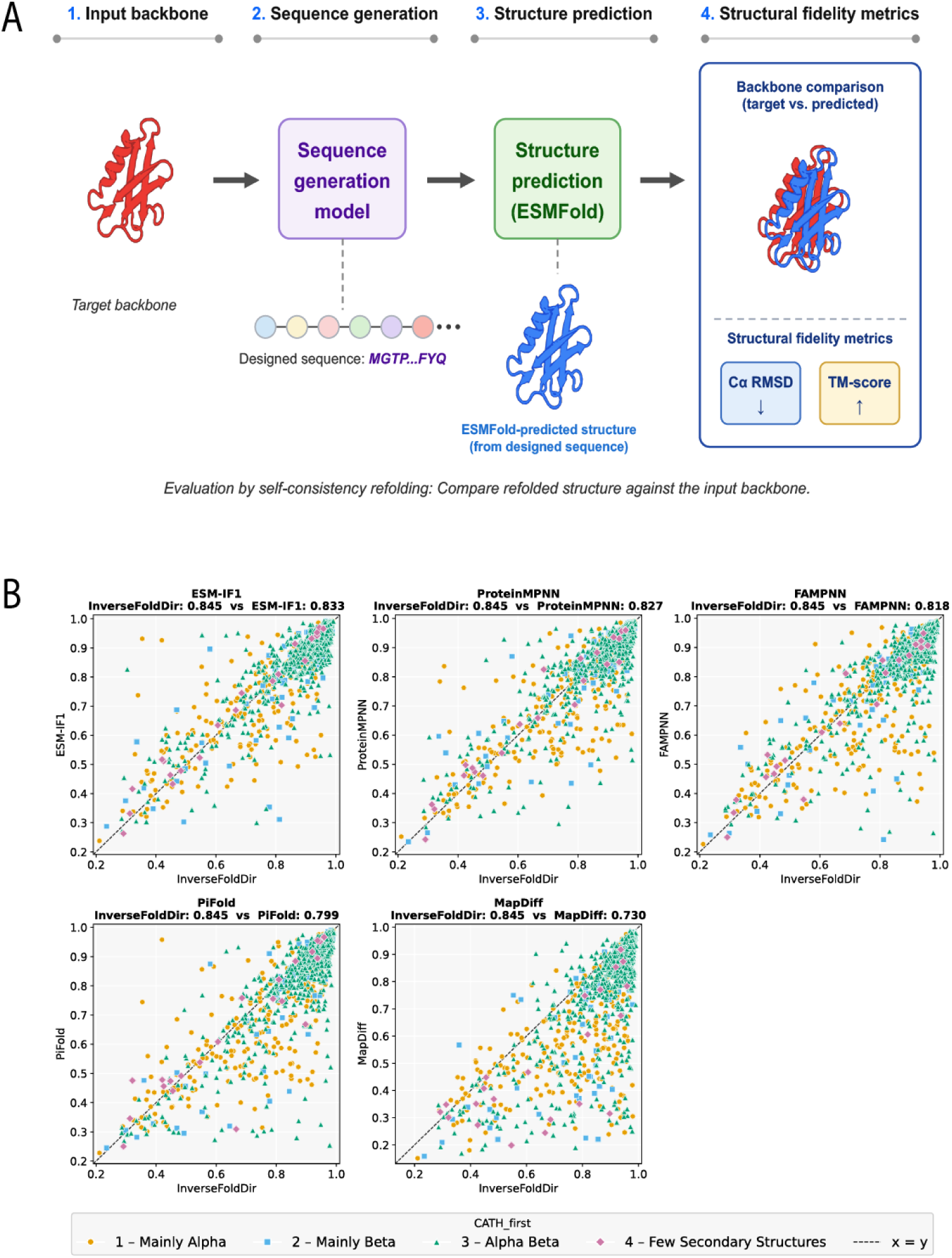
Evaluation of predicted structural recovery. (A) The evaluation pipeline for the predicted structural recovery of Inverse FoldDir and competing models uses ESMFold as the structure predictor. (B) Per-protein comparisons between Inverse FoldDir and competing methods, with each point representing the mean TM-score over three replicates. Points below the diagonal indicate proteins for which Inverse FoldDir outperforms the competing method, whereas points above the diagonal indicate proteins for which the competing method outperforms Inverse FoldDir.

Across the benchmark, most proteins clustered near or under the diagonal when compared against the competing methods on a per-protein basis, indicating that the improvement was broadly distributed rather than driven by a small number of favorable examples (Figure 4B).

On an aggregate level, Inverse FoldDir achieved the highest structural self-consistency among the evaluated methods. It produced the highest average TM-score [15] and the lowest average C*α* RMSD, outperforming ESM IF [2], ProteinMPNN [6], FAMPNN [9], PiFold [10], and MapDiff [16] under the specified inference settings and shared evaluation pipeline. Pairwise statistical tests demonstrated statistically significant performance gains in the predicted structural recovery metrics (Table 1).

**Table 1:** Structure-recovery benchmark with pairwise *t*-test significance relative to the strongest-performing method for each metric. Values are means across 1,120 proteins, with standard deviations across three generation-replicate means, each averaged over the complete test set. Tests used 1,120 per-protein means per method, each calculated across the three generation replicates.

| Model | Approach | TM-score ↑ | RMSD ($\text{\AA}$) ↓ |
| --- | --- | --- | --- |
| ESM-IF1 | Autoregressive | 83.3±0.2*** | 1.86±0.01*** |
| ProteinMPNN | Autoregressive, random unmasking | 82.7±0.1*** | 1.89±0.01*** |
| FAMPNN | Autoregressive + coordinate diffusion | 81.8±0.3*** | 1.95±0.02*** |
| PiFold | One shot | 79.9±0.1*** | 2.06±0.01*** |
| MapDiff | Discrete Denoising Diffusion + IPA | 73.0±0.3*** | 2.52±0.02*** |
| Inverse FoldDir (ours) | Dirichlet Denoising Flow Matching | **84.5±0.2** | **1.76±0.01** |

\*, \*\*, and \*\*\* denote a statistically significant difference from the top performer (bold) at $p < 0.05$ , $p < 0.01$ , and $p < 0.001$ , respectively, based on a one-tailed pairwise $t$ -test with the alternative hypothesis that the top performer is superior in the favorable direction for the metric (higher TM-score or lower RMSD).

We next examined performance across the major CATH secondary-structure classes (Table 2). Inverse FoldDir maintained superior structural recovery across mainly *α*, mainly *β*, and *α*-*β* proteins but failed to improve in the “few secondary structures” category.

**Table 2:**
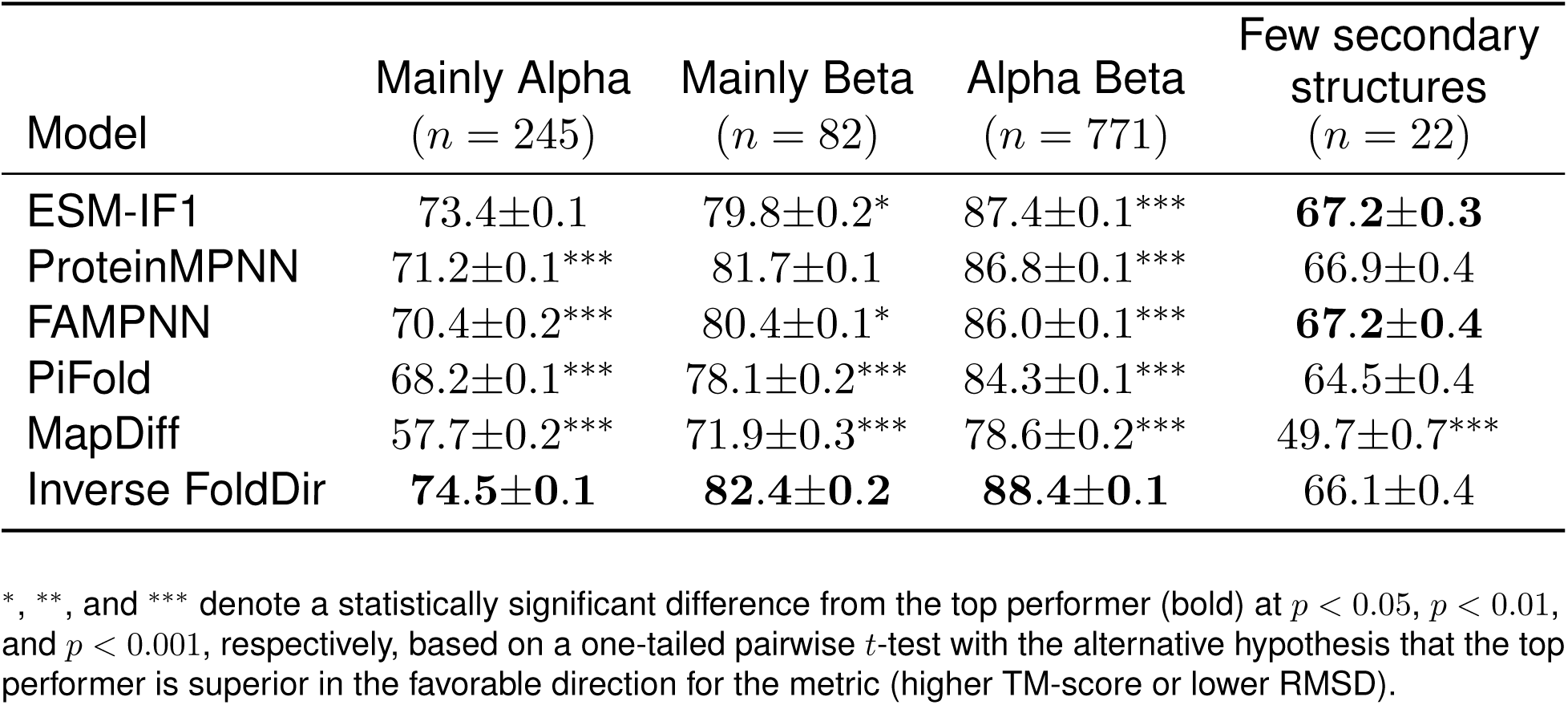
Structure-recovery TM-scores on the test set by CATH secondary-structure class.

| Model | Mainly Alpha<br>($n = 245$) | Mainly Beta<br>($n = 82$) | Alpha Beta<br>($n = 771$) | Few secondary<br>structures<br>($n = 22$) |
| --- | --- | --- | --- | --- |
| ESM-IF1 | 73.4±0.1 | 79.8±0.2* | 87.4±0.1*** | **67.2±0.3** |
| ProteinMPNN | 71.2±0.1*** | 81.7±0.1 | 86.8±0.1*** | 66.9±0.4 |
| FAMPNN | 70.4±0.2*** | 80.4±0.1* | 86.0±0.1*** | **67.2±0.4** |
| PiFold | 68.2±0.1*** | 78.1±0.2*** | 84.3±0.1*** | 64.5±0.4 |
| MapDiff | 57.7±0.2*** | 71.9±0.3*** | 78.6±0.2*** | 49.7±0.7*** |
| Inverse FoldDir | **74.5±0.1** | **82.4±0.2** | **88.4±0.1** | 66.1±0.4 |

\*, \*\*, and \*\*\* denote a statistically significant difference from the top performer (bold) at $p < 0.05$ , $p < 0.01$ , and $p < 0.001$ , respectively, based on a one-tailed pairwise $t$ -test with the alternative hypothesis that the top performer is superior in the favorable direction for the metric (higher TM-score or lower RMSD).

The ablation study (Table 3; Supplementary Section S8) showed that removing the additional AlphaFold-predicted training structures or the iterative denoising updates significantly reduced structural recovery, whereas removing auxiliary components such as the multitask prediction heads or mixed-mode training produced comparatively modest changes.

**Table 3:** Ablation experiments on the CATH 4.2 test set.

| Model version | TM-score ↑ | RMSD ($\text{\AA}$) ↓ |
| --- | --- | --- |
| Base model | **84.5±0.2** | **1.76±0.01** |
| Removed: AlphaFold predicted structures | 81.0±0.2*** | 2.02±0.01*** |
| Removed: Iterative denoising updates | 83.6±0.2*** | 1.87±0.01*** |
| Removed: Multitasking heads (charge, steric) | 84.2±0.1 | 1.78±0.01 |
| Removed: Mixed mode training | 84.3±0.1 | 1.78±0.01 |
| Removed: Entropy | 84.3±0.2 | 1.78±0.02 |
| Removed: Burial descriptors | 84.3±0.1 | 1.78±0.01 |

\*, \*\*, and \*\*\* denote a statistically significant difference from the top performer (bold) at $p < 0.05$ , $p < 0.01$ , and $p < 0.001$ , respectively, based on a one-tailed pairwise $t$ -test with the alternative hypothesis that the top performer is superior in the favorable direction for the metric (higher TM-score or lower RMSD).

### 3.4 Inverse FoldDir likelihoods capture structure-linked mutational tolerance on ProteinGym

We tested whether Inverse FoldDir could provide information about the effects of individual aminoacid substitutions beyond sequence generation. For each position, we computed log-likelihood ratios by comparing the probabilities assigned to mutant and wild-type residues using a position-wise inpainting procedure and evaluated these scores on 35 deep mutational scanning assays from the ProteinGym database [17] (Supplementary Data 1).

Across a range of log-likelihood-ratio thresholds, Inverse FoldDir was competitive with the strongest model in zero-shot positive predictive value (Figure 5A). Prediction performance depended strongly on assay type. Inverse FoldDir likelihood ratios showed the strongest agreement with stability-related assays, whereas performance was lower for assays measuring binding affinity, enzymatic activity, or other functional phenotypes. The performance differences were not statistically significant (Figure 5B and Supplementary Figure S3).

**Figure 5:**
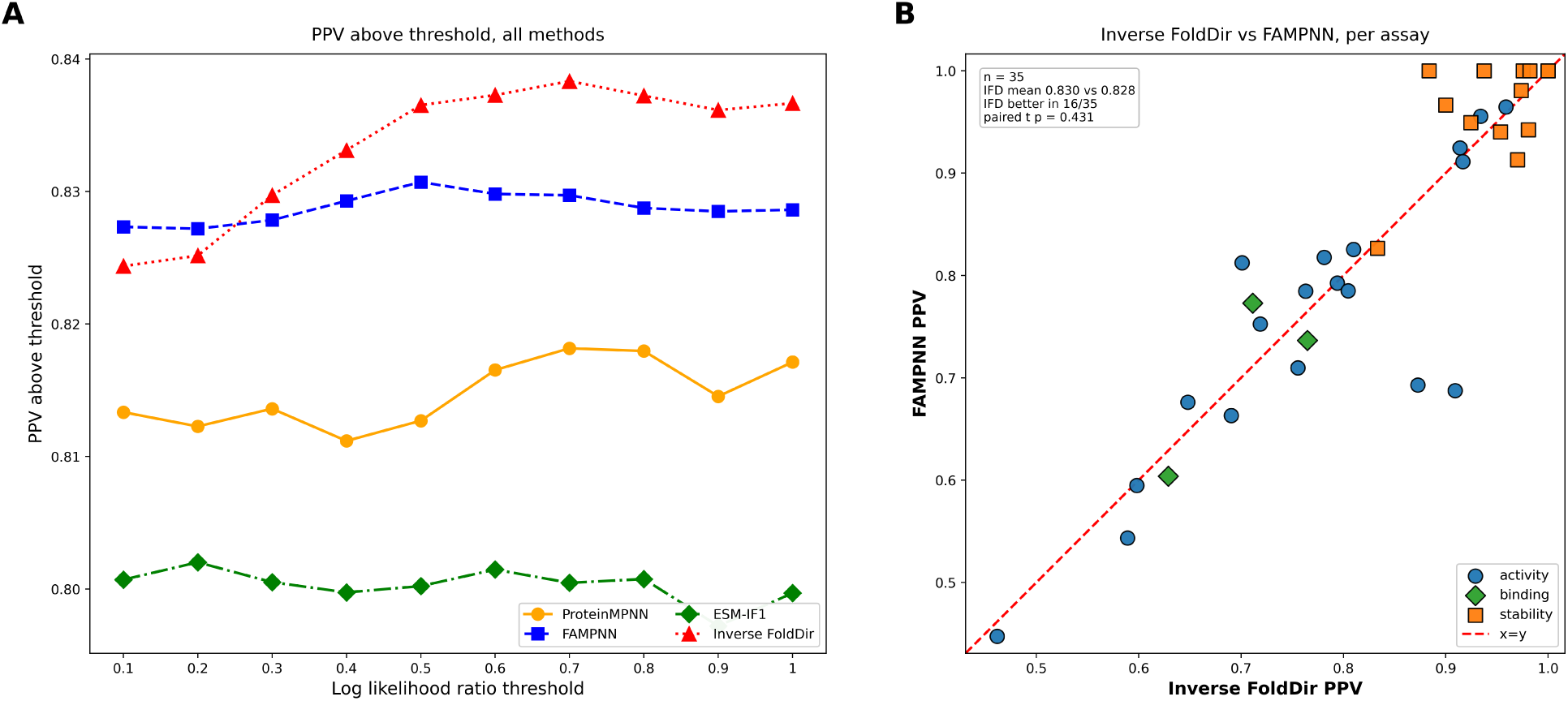
ProteinGym positive predictive value (PPV) performance. (A) PPV above increasing log-likelihood-ratio thresholds for Inverse FoldDir and inverse-folding baselines. (B) Per-assay comparison of Inverse FoldDir and FAMPNN PPV, grouped by assay phenotype.

### 3.5 Inverse FoldDir redesigns anti-GFP nanobody sequences that retain binding

We asked whether structural self-consistency could support the redesign of a functional sequence from an experimentally validated protein scaffold under single-chain conditioning. As a representative test case, we selected an X-ray-crystallized anti-GFP nanobody (PDB: 3OGO) [18] that was not present in our training set and evaluated whether Inverse FoldDir could generate substantially different nanobody sequences that retained measurable GFP binding as a proxy for structural fidelity. Excluding the His tag, the nanobody contains 116 residues, of which 105 have resolved coordinates.

We performed sequence generation using only the nanobody backbone as structural input, without providing GFP during generation (Figure 6A). This single-chain setting tests whether the model can redesign a protein while preserving its overall structural framework rather than relying on explicit information about the binding partner.

**Figure 6:**
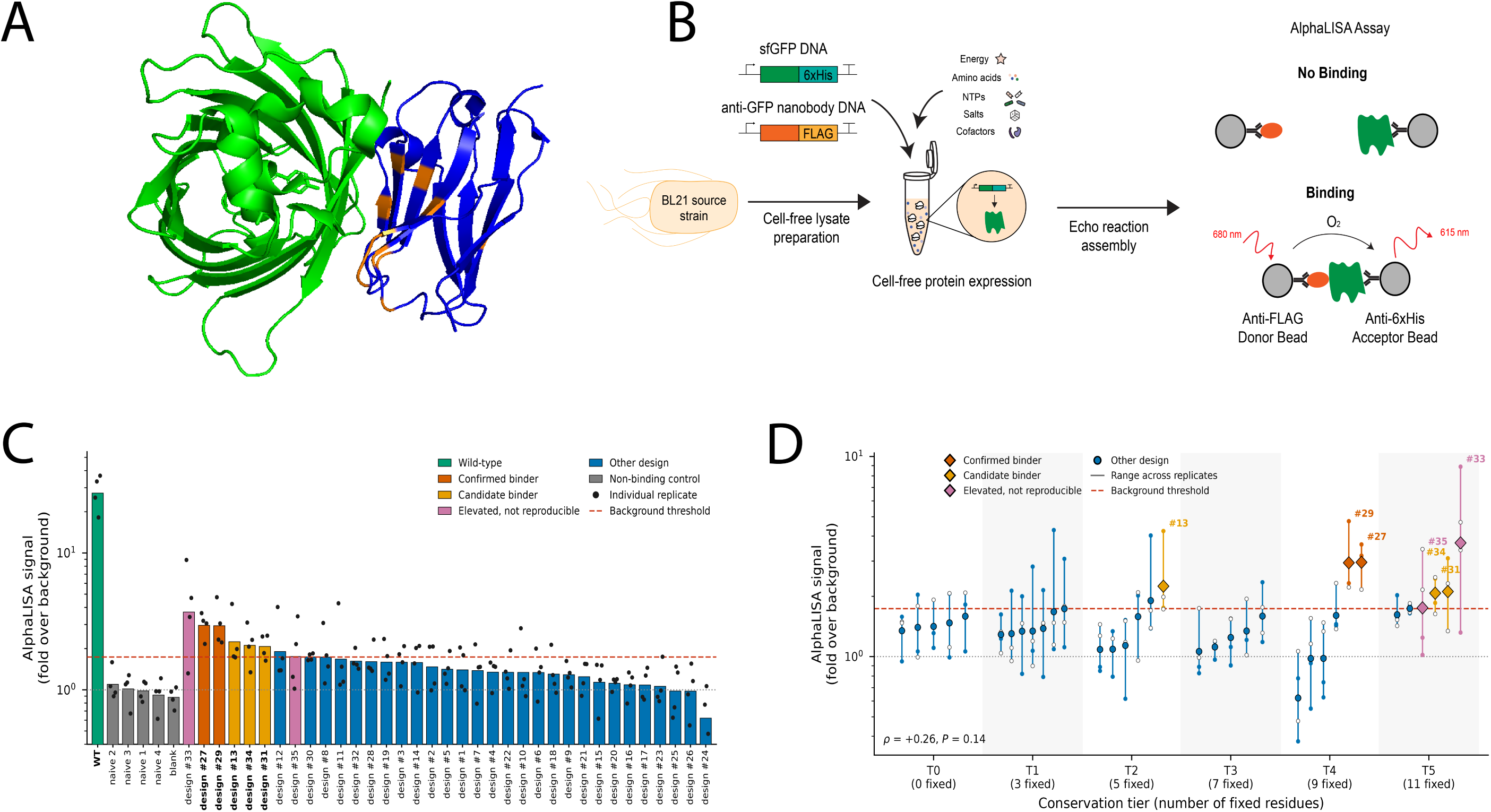
Experimental validation of redesigned anti-GFP nanobodies. (A) Structure of the nanobody-GFP complex (PDB: 3OGO), with GFP shown in green, the nanobody in blue, and positions selected for fixation based on hypothesized roles in structural integrity or binding highlighted in orange. (B) Experimental workflow. Thirty-five Inverse FoldDir designs, the wild-type (WT) nanobody, and four naive negative-control sequences were expressed in a cell-free system, and binding to sfGFP-6×His was measured using a luminescent AlphaLISA assay. (C) AlphaLISA signal expressed as fold over the five controls (four naive-baseline sequences and a no-template blank) measured in the corresponding replicate block. Bars show the geometric mean of four measurements, and dots show individual measurements. The background threshold was defined as the control mean plus three standard deviations using variance pooled across all 20 control observations (1.74-fold). The dotted line indicates the control mean. Confirmed binders exceeded the threshold in all four measurements, whereas candidate binders exceeded it in three of four measurements. (D) Designs grouped by conservation tier, from T0 (no fixed positions) to T5 (11 fixed positions). Vertical lines span the minimum and maximum of the four replicate-matched measurements. Small filled and open markers denote individual measurements from runs 1 and 2, respectively, and large markers denote geometric means.

We experimentally tested 40 constructs in total: the wild-type nanobody as a positive control, 35 Inverse FoldDir designs, and 4 naive-baseline sequences. The designs spanned sequence-constraint levels ranging from 0 to 11 fixed residues among 105 modeled positions, selected a priori as conservative safeguards for structural integrity or binding (Supplementary Section S2). C23 and C97 were recovered in every generated Tier 0 sequence, showing that their native identities could be preserved without explicit fixation. The naive-baseline sequences were created through random substitutions without guidance from a structural or generative AI model. These baseline sequences had 55% sequence identity and 75% to 100% sequence similarity to the native sequence, with predicted structural self-consistency TM-scores ranging from 72.0 to 97.6 on the 0-100 reporting scale. Complete design sequences, tiers, and associated computational metadata are provided in Supplementary Data 2.

We experimentally evaluated 35 Inverse FoldDir designs using an anti-GFP nanobody scaffold. All constructs were expressed in a cell-free system and assayed for binding to sfGFP-6×His using AlphaLISA [19]. Binding was measured in two independent experiments and normalized to non-binding controls measured in the same assay. Two designs, #27 and #29, showed reproducible binding above the background threshold across all measurements and were classified as confirmed binders (Figure 6B-D). Three additional designs, #13, #31, and #34, showed elevated binding in three of four measurements and were considered candidate binders. The wild-type nanobody produced a substantially stronger signal than all redesigned sequences. Two additional designs, #33 and #35, showed high mean signals but substantial variability between replicates and were therefore not classified as reproducible binders.

Both confirmed binders were generated in Tier 4, in which only 9 of 105 modeled positions were fixed. Design #27 retained 57% sequence identity to the native nanobody while producing a geometric-mean AlphaLISA signal of 2.96-fold over background. Design #29 produced a comparable 2.94-fold signal. Thus, Inverse FoldDir generated substantially redesigned sequences that retained reproducible antigen-binding activity.

As an orthogonal computational control, Boltz-2 [20] cofolding confidence did not correlate with AlphaLISA signal either with an MSA (Spearman *ρ* = 0.06, *P* = 0.74) or in single-sequence mode (*ρ* = −0.03, *P* = 0.86; Supplementary Figure S4; analysis details in Supplementary Section S7.5). Other evaluated structure-confidence metrics likewise showed no significant association with binding signal (Supplementary Figure S5), and ROC analyses did not support using these metrics to rank the two confirmed binders (Supplementary Figure S6).

## 4 Discussion

### 4.1 Inverse FoldDir improves structural recovery across diverse protein backbones

We introduced Inverse FoldDir, a controllable method for protein inverse folding that performs iterative denoising over full-sequence amino-acid probability distributions. The method addresses two related design settings: (1) redesigning an entire sequence or selected regions of an existing protein scaffold and (2) assigning structurally plausible sequences to computationally generated backbones for which no natural sequence exists. Inverse FoldDir supports complete sequence generation, fixed-residue inpainting, and user-defined soft residue priors within the same generative framework. Although predicted structural recovery is an imperfect proxy for downstream experimental success, it measures compatibility with the target backbone more directly than native sequence recovery because many distinct sequences can adopt the same fold [6, 2]. We therefore use predicted structural self-consistency as our primary measure of inverse-folding performance. Across held-out CATH 4.2 backbones [1], Inverse FoldDir achieved greater predicted structural self-consistency than the evaluated inverse-folding baselines. These gains extended across three of the four major CATH classes rather than being confined to a single structural category. Results for proteins in the “few secondary structures” class should be interpreted cautiously because this subset contained only 22 test proteins and therefore exhibited greater variability than the larger classes. This category also contains a higher prevalence of unstructured and flexible regions, making structural recovery more challenging than in classes containing more stable secondary structures. The improvement from AlphaFoldDB augmentation [21] (Table 3) suggests that inverse-folding models benefit from broader exposure to protein structural space. We used the published AlphaFoldDB cluster organization [22] to construct a confidence-filtered training corpus with broad structural coverage while controlling redundancy and preserving separation from the held-out CATH benchmark (Supplementary Section S4). Because structurally similar clusters were explicitly excluded, the resulting performance gain is consistent with broader training-set coverage contributing to generalization rather than simply increasing exposure to structures closely related to the benchmark.

Together, these results establish Dirichlet flow matching [11] as an effective framework for structure-conditioned protein sequence generation and demonstrate its practical use in binding-preserving scaffold redesign. They do not, however, imply that every structurally self-consistent sequence will express efficiently, remain soluble, fold robustly, or retain a desired biological function. Structural recovery is therefore best viewed as a necessary but insufficient design criterion: poor recovery can eliminate unsuitable candidates, whereas strong recovery alone does not ensure experimental activity. Practical protein design requires structural compatibility to be considered alongside objectives such as stability, solubility, expression, and function.

### 4.2 Iterative denoising enables whole-sequence refinement and flexible design control

The iterative denoising formulation provides a design interface that differs fundamentally from one-shot and autoregressive sequence generation [2, 6, 10]. Inverse FoldDir maintains a mutable probability distribution at every nonfixed position throughout inference. Residues resolve at different rates, and some positions revise their most likely amino-acid identity after neighboring positions have begun to converge. The resulting trajectory portrays sequence design as coordinated whole-sequence refinement rather than a series of independent or irreversible assignments. Autoregressive models often generate protein sequences that have high structural fidelity, and our trajectory analysis is not intended to establish that iterative denoising universally outperforms sequential decoding. Instead, it demonstrates a distinct capability: Inverse FoldDir can revisit earlier sequence preferences as broader sequence context develops, while also exposing position-level uncertainty and commitment during generation. We also show that iterative updating improves structural recovery relative to predicting the entire sequence in a single step.

A central aim of Inverse FoldDir is to let protein designers express different levels of prior knowledge. Fixed-residue inpainting preserves exact identities at positions such as catalytic residues, disulfide-forming cysteines, sequence motifs, or known interaction hot spots. Soft residue priors provide a less restrictive form of control: users can bias selected positions toward residue classes or custom amino-acid distributions without committing them to one identity. This capability may prove useful when a designer knows the desired biochemical character of a region but does not know the optimal sequence, such as favoring polar residues on an exposed surface or allowing several possible metal-coordinating residues. The continuous denoising state also provides a natural interface for property-guided sampling. When differentiable scoring functions exist, future implementations could steer generation toward objectives such as net charge, solubility, aggregation propensity, expression, stability, or predicted interface compatibility during sampling rather than filtering only after sequence generation. The present study experimentally evaluates full generation and fixed-residue inpainting, implements soft-prior conditioning, and leaves systematic property-guided optimization for future work.

### 4.3 Structural conditioning supports protein redesign but does not fully determine biological function

The anti-GFP nanobody experiment tested whether computational structural recovery could translate into retention of an experimentally measured function. Inverse FoldDir generated candidate sequences using only the nanobody backbone, without providing GFP as context during generation. Several selected designs retained measurable GFP-binding signal despite more than 40% sequence divergence from the native nanobody sequence, including designs produced with 9 of 105 modeled positions fixed. This result provides experimental support for binding-preserving scaffold redesign and strengthens the interpretation of the CATH 4.2 benchmark [1]: sequences that recover an intended backbone computationally can, in at least one experimentally tested system, also retain a biological phenotype. However, the lack of an association between Boltz-2 [20] confidence and experimental signal further shows that predicted structural or complex plausibility is insufficient to establish binding. We deliberately interpret this experiment as the redesign of a known binding scaffold rather than de novo binder discovery. The input backbone came from a validated anti-GFP nanobody (PDB: 3OGO) [18], and some design tiers incorporated prior knowledge about residues associated with structural stability or binding.

The ProteinGym analysis [17] further clarified what information the model learns. Position-wise likelihood ratios showed the strongest utility in stability-related deep mutational scanning assays and weaker alignment with binding or activity measurements. This pattern matches the model objective: Inverse FoldDir learns which sequences and substitutions remain compatible with a single-chain protein backbone, but it does not directly model biochemical phenotypes or interaction partners. We focused primarily on positive predictive value because protein-design workflows often prioritize a limited set of top-ranked mutations for experimental testing rather than requiring an accurate global ordering of all variants. Many mutations that disrupt binding-interface chemistry are less disruptive to the global fold, causing these binding-disruptive mutations to be interpreted as tolerated from the perspective of global folding. Inverse FoldDir likelihoods contain information about structure-linked mutational tolerance, particularly for stability-like phenotypes, but the method should not be treated as a general-purpose variant-effect predictor without additional functional objectives, biological context, or task-specific calibration.

Several limitations define the scope of the present study. First, the experimental validation was limited to one anti-GFP nanobody system. Additional testing across protein families, folds, and functional assays will determine how broadly the observed binding preservation generalizes.

Second, AlphaLISA measures relative binding signal rather than structural recovery, whereas the model was designed to recover backbone structure. Third, Inverse FoldDir was trained primarily on single-chain structures and does not yet constitute a general protein-complex design model. Fourth, the study implements soft residue priors but does not systematically measure how strongly they shift residue composition, preserve structural recovery, or improve experimental outcomes. Likewise, although the continuous trajectory enables future property-guided sampling in principle, we do not demonstrate general multi-objective optimization here.

Despite these limitations, Inverse FoldDir establishes a general strategy for controllable protein inverse folding. The method combines superior predicted structural recovery with an interpretable, mutable generation trajectory and supports multiple forms of user input within one representation. Experimental nanobody redesign shows that this framework can produce substantially altered sequences that retain measurable binding. Future work can extend the framework to multichain complexes, integrate experimentally calibrated property objectives, and couple generation with iterative design-build-test cycles. Inverse FoldDir therefore provides both a validated inverse-folding method and a foundation for more controllable, property-aware protein sequence design.

## 5 Materials and methods

### 5.1 Model formulation

We formulated inverse folding as the conditional generation of an amino-acid sequence from a protein backbone. For the amino-acid objective, we followed the Dirichlet flow-matching framework of Stark et al. [11], which represents categorical sequences as continuous probability distributions on the simplex and learns a vector field that transports uncertain distributions toward categorical sequence states. We adapted this framework to protein sequences by representing each residue as a probability vector over 21 classes (the 20 canonical amino acids and an unknown-residue class) and conditioning the amino-acid predictions on an SE(3)-equivariant residue-graph representation of the input backbone. The probability path, structure-conditioned vector field, numerical integration procedure, and mathematical treatment of fixed residues and soft priors are detailed in Supplementary Section S1.

During training, we sampled intermediate sequence states from the time-dependent Dirichlet probability path defined by Stark et al. [11] and trained the network to predict the native amino-acid identity at each nonfixed position. The complete training procedure is summarized in Supplementary Algorithm 2. During inference, we independently initialized residue distributions from an uninformative Dirichlet distribution and numerically integrated the learned structure-conditioned flow for 20 steps. We projected residue vectors onto the simplex after each update and decoded the final sequence by selecting the highest-probability amino acid at each position.

For fixed-residue inpainting, we held user-specified amino-acid identities constant throughout integration and updated only the remaining positions. For soft-prior conditioning, we replaced the default initial distribution at selected positions with user-defined amino-acid probability distributions while allowing those positions to change during denoising. Detailed graph construction, feature definitions, and sampling parameters are provided in Supplementary Sections S3 and S5.

### 5.2 Protein graph construction

We represented each protein backbone as a residue-level geometric graph and encoded it using an SE(3)-equivariant network based on the Geometric Vector Perceptron (GVP) architecture [23]. Edges connected residues based on spatial proximity, using a 12 Å C*α* distance cutoff, rather than purely on sequence adjacency. The graph combined backbone geometry with the current amino-acid probability state, allowing structural context and evolving sequence uncertainty to be processed jointly during denoising. Complete graph-construction procedures, feature definitions, calculations, and dimensions are provided in Supplementary Section S3 and Supplementary Tables S1, S2, and S3.

### 5.3 Model architecture

Inverse FoldDir combines the Dirichlet flow-matching framework of Stark et al. [11] with an SE(3)-equivariant graph neural network based on the Geometric Vector Perceptron (GVP) architecture of Jing et al. [23]. At each denoising step, the model receives the residue graph together with the current amino-acid probability distributions and a learned embedding of the denoising time. A stack of GVP graph-attention layers [23] propagates information across the protein while jointly updating scalar and vector node representations, allowing residue identities to evolve in the context of both local geometry and neighboring sequence uncertainty.

A prediction head maps the final node representations to a categorical distribution over the 21 amino-acid classes at every residue position. During training, the model learns to predict the native amino-acid identities from intermediate Dirichlet samples generated along the probability path. In addition to the primary sequence prediction head, we jointly optimized auxiliary charge and steric prediction heads to provide additional physicochemical supervision during training so that the model receives partial credit for predicting a different amino acid with similar geometric or electrostatic properties. These auxiliary tasks contribute only to the training objective and are not required during inference.

Inverse FoldDir contains 9.7 million trainable parameters. Complete optimization settings, computational resource details, and model hyperparameters are reported in Supplementary Section S5, and the auxiliary prediction tasks are detailed in Supplementary Section S6 and Supplementary Table S4.

### 5.4 Training data

Inverse FoldDir was trained using the CATH 4.2 training split [1] together with a structurally filtered set of AlphaFoldDB structures [21]. To expand structural coverage while limiting redundancy, we constructed the AlphaFoldDB training corpus using the structural-clustering framework of Barrio-Hernandez et al. [22]. Candidate clusters were filtered for structural confidence and separation from the held-out CATH 4.2 validation and test sets before expansion to individual structures. Training clusters whose representatives showed Foldseek similarity to a held-out representative at *E <* 0.1 were excluded in their entirety, removing all members of each identified cluster [24]. We additionally verified that residual structural similarity introduced by the retained AlphaFoldDB clusters was no greater than the range already present between the standard CATH 4.2 training and held-out sets. Experimental and predicted structures were sampled jointly during training. Complete dataset construction and filtering criteria are provided in Supplementary Section S4.

### 5.5 Sequence generation and sampling

For sequence generation, unconstrained residues were initialized from an uninformative Dirichlet distribution, fixed residues were clamped to one-hot identities, and soft priors replaced the initial distributions at selected positions. We integrated the structure-conditioned flow for 20 denoising steps and decoded the terminal state by selecting the highest-probability amino acid at each residue. Algorithm 1 summarizes generation. Complete numerical and sampling details are provided in Supplementary Sections S1 and S5.

In Algorithm 1, *s* denotes normalized integration progress and *t* = 100*s* is the Dirichlet time used for model conditioning and flow integration. The sampling temperature *τ* is applied to amino-acid logits before softmax to obtain the probabilities used to construct the flow. We used *τ* = 1, leaving the logits and resulting probabilities unchanged. This sampling parameter is not applied to training or final argmax decoding.

#### Algorithm 1 Structure-conditioned sequence generation with Inverse FoldDir

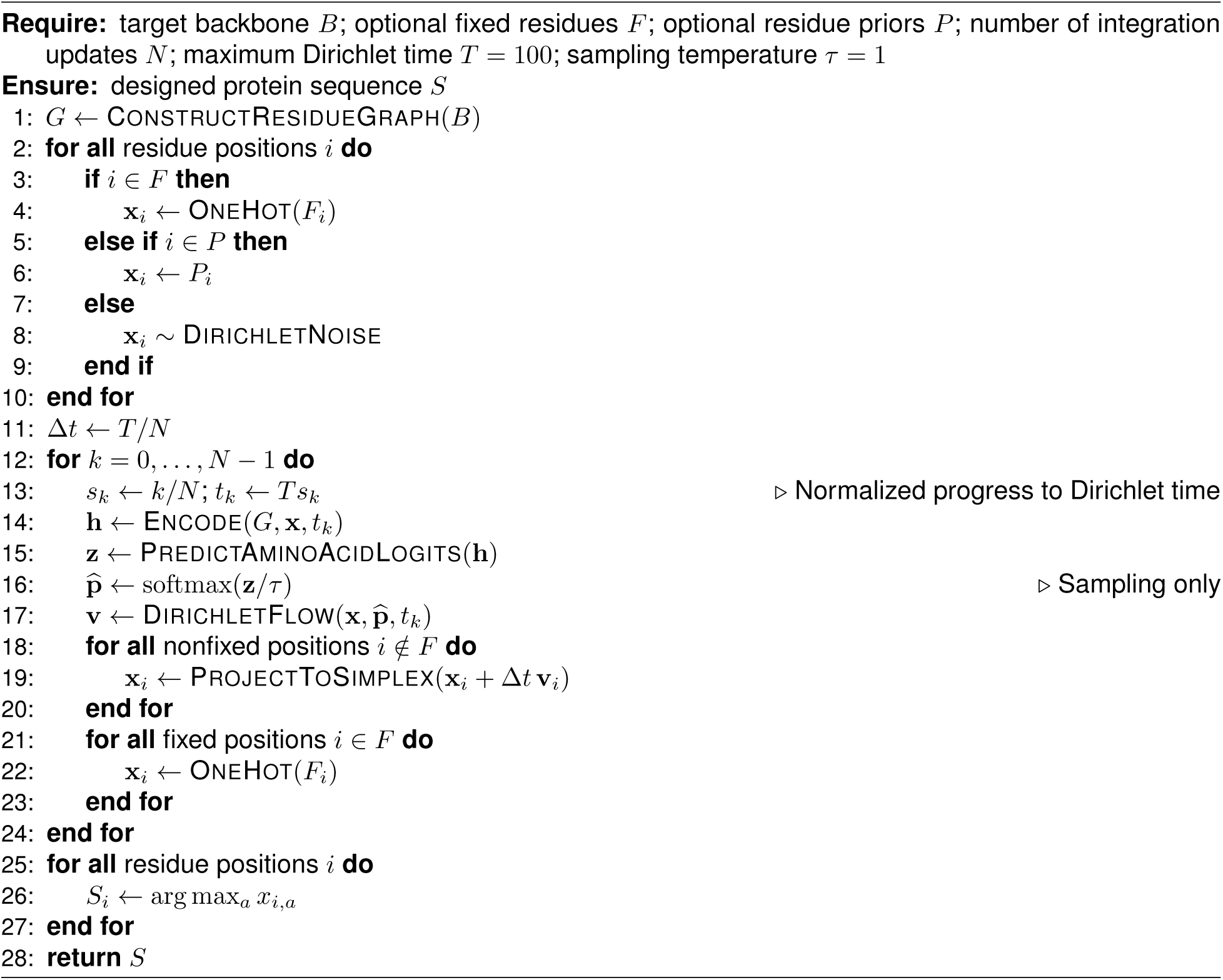

### 5.6 Cell-free protein expression

Cell-free extracts were generated following previously reported protocols [25, 26, 27]. Anti-GFP nanobody designs, negative controls, and sfGFP-6×His were expressed using an *E. coli* BL21(DE3*) PANOx-SP cell-free expression system [28] under oxidizing conditions based on prior cell-free production of disulfide-bonded proteins [29]. These conditions were used to promote nanobody disulfide-bond formation. The complete cell-free expression protocol is provided in Supplementary Section S7.

### 5.7 AlphaLISA binding assay

Binding between cell-free-expressed nanobodies and sfGFP-6×His was quantified using an AlphaLISA proximity assay [19]. The workflow was adapted from previous cell-free expression and binding-screening studies [30, 31] and optimized using the wild-type nanobody, after which redesigned variants were evaluated under identical conditions. Complete experimental procedures are provided in Supplementary Section S7.

AlphaLISA signals were normalized to the five controls (one blank and four naive baselines) measured under the corresponding assay condition. A background threshold of three standard deviations above the pooled non-binding-control signal was used for classification. Designs exceeding this threshold in all four analyzed measurements were classified as confirmed binders, whereas designs exceeding the threshold in three of four measurements were classified as candidate binders. Additional analysis details are provided in Supplementary Section S7.4.

### 5.8 ProteinGym analysis

To assess whether Inverse FoldDir likelihoods capture information related to mutational tolerance, we evaluated the model on 35 deep mutational scanning datasets from ProteinGym [17]. All ProteinGym evaluations were restricted to single-amino-acid substitution variants. For each mutation, a position-wise inpainting procedure was used to estimate the log-likelihood ratio between the mutant and wild-type amino acids while conditioning on the native backbone structure. Experimental positives for positive predictive value calculations were defined using the assay-specific binary labels in ProteinGym’s preprocessed tables. Complete baseline configurations, scoring protocols, dataset composition, and evaluation details are provided in Supplementary Section S9 and Supplementary Table S5.

## 6 Data availability

Software, pretrained model weights, documentation, and example notebooks for full-sequence generation, fixed-residue inpainting, soft-prior initialization, and candidate refolding and ranking are available at github.com/AlpTartici/inversefolddir. Zero-shot ProteinGym results will be provided as Supplementary Data 1, and complete nanobody design sequences, constraint tiers, processed experimental results, and associated computational metadata will be provided as Supplementary Data 2. These supplementary datasets will be uploaded as separate files upon publication.

## 7 Author contributions

A. Tartici and B.J.W. conceived the project and developed the initial methodology and computational framework. A. Tartici implemented, extended, and evaluated Inverse FoldDir, including its open-source implementation, and performed the computational experiments and analyses, with contributions from M.S. to software development, experiments, and analysis. A. Tartici and A. Tian designed the experimental validation studies, which A. Tian performed and analyzed under the supervision of M.C.J. B.J.W. supervised the initial development, and R.B.A. supervised the subsequent development and evaluation and provided computational resources. A. Tartici wrote the original manuscript with input from the other authors. All authors reviewed and edited the manuscript. R.B.A. and B.J.W. jointly supervised the work.

## 8 Acknowledgments

This work was supported by the Army Research Office (W911NF-22-2-0246), the National Science Foundation (TIP-2448820), NIH grant GM153195, and Stanford Medical Scholars support. R.B.A. is supported by NIH grants GM102365 and AG089509 and Burroughs Wellcome Fund award 1074128. Alp Tartici acknowledges support from the Stanford Graduate Fellowship (Smith Fellowship). Anru Tian acknowledges support from the National Science Foundation Graduate Research Fellowship under grant no. DGE-2146755. The Microsoft Office of the Chief Scientific Officer supported this research and Alp Tartici during his internship there. We thank Dr. Stephen Montgomery, Dr. Brian Hie, Dr. Anshul Kundaje, and the members of the Altman laboratory for useful discussions. We thank Stanford Research Computing for providing computational resources that contributed to this research.

