## Supplemental File for "Inverse FoldDir: Structure-conditioned Protein Sequence Design by Dirichlet Flow Matching"

**S10 Supplementary figures** **32**

### S1 Mathematical formulation of structure-conditioned Dirichlet flow matching

#### S1.1 Probability-simplex representation of protein sequences

Inverse FoldDir adapts Dirichlet flow matching [1] to structure-conditioned protein sequence generation. Related simplex-based generative approaches have represented categorical biological sequences as continuous variables on a probability simplex, including the Dirichlet diffusion score model of Avdeyev et al. [2]. Whereas that approach defines a stochastic diffusion process with a Dirichlet stationary distribution, Inverse FoldDir follows the deterministic Dirichlet flow-matching formulation of Stark et al. and conditions the generative process on protein backbone geometry.

For a categorical variable with $K$ possible states, define the probability simplex

$$\Delta^{K-1} = \left\{ \mathbf{x} \in \mathbb{R}^K : x_a \geq 0,\ \sum_{a=1}^{K} x_a = 1 \right\}. \tag{S1}$$

Inverse FoldDir uses $K = 21$ model classes, corresponding to the 20 canonical amino acids and an additional unknown or masked class. A protein of length $L$ is represented during generation by

$$\mathbf{X}_t = (\mathbf{x}_{1,t}, \ldots, \mathbf{x}_{L,t}) \in \left(\Delta^{K-1}\right)^L, \tag{S2}$$

where $\mathbf{x}_{i,t}$ is the amino-acid probability vector at residue $i$ at denoising time $t$. A discrete amino-acid identity $y_i$ corresponds to a vertex of the simplex represented by the one-hot vector $\mathbf{e}_{y_i}$.

This continuous representation permits uncertainty over amino acid identity to be retained during generation. Rather than assigning a residue once and treating that assignment as irreversible, all designable positions can remain within the interiors of their respective simplices and can therefore change their most probable amino-acid identities as generation proceeds.

#### S1.2 Dirichlet probability path

Following Stark et al. [1], the transition between an uninformative simplex state and a categorical amino-acid state is defined using a family of Dirichlet distributions. The Dirichlet density for parameter vector $\boldsymbol{\alpha}$ is

$$\mathrm{Dir}(\mathbf{x};\boldsymbol{\alpha}) = \frac{1}{B(\boldsymbol{\alpha})}\prod_{a=1}^{K} x_a^{\alpha_a - 1}, \tag{S3}$$

where $B(\boldsymbol{\alpha})$ is the multivariate beta function. For a residue whose target identity is $y_i$, the conditional probability path is

$$p_t\left(\mathbf{x}_{i,t} \mid y_i\right) = \mathrm{Dir}\left(\mathbf{x}_{i,t}; \mathbf{1} + t\mathbf{e}_{y_i}\right). \tag{S4}$$

At $t = 0$, this reduces to the uniform Dirichlet distribution,

$$p_0(\mathbf{x}_{i,0}) = \mathrm{Dir}(\mathbf{x}_{i,0}; \mathbf{1}), \tag{S5}$$

which has support throughout the simplex and does not favor any amino-acid class. As $t$ increases, probability mass progressively concentrates toward the vertex $\mathbf{e}_{y_i}$. In practice, the flow is integrated to a finite maximum time rather than to the limiting distribution, after which the terminal probability vector is discretized.

For a protein sequence, intermediate states are sampled independently across positions conditional on the native sequence,

$$p_t(\mathbf{X}_t \mid \mathbf{y}) = \prod_{i=1}^{L} p_t(\mathbf{x}_{i,t} \mid y_i). \tag{S6}$$

Although the corruption process happens independently across residue positions, the denoising model does not. Inverse FoldDir jointly processes all residue states together with the three-dimensional protein structure, allowing the prediction at one position to depend on the structural and evolving sequence context of other residues.

#### S1.3 Structure-conditioned denoising model

Let $\mathcal{G}(\mathbf{B})$ denote the residue-level geometric graph constructed from target backbone coordinates $\mathbf{B}$, as detailed in Supplementary Section S3. At time $t$, the neural network receives the graph, the complete noisy sequence state $\mathbf{X}_t$, and a learned representation of $t$, and predicts a categorical distribution over amino-acid identities at every residue:

$$\widehat{p}_{\theta,i}\left(a \mid \mathbf{X}_t, \mathcal{G}(\mathbf{B}), t\right), \qquad a \in \{1, \ldots, K\}. \tag{S7}$$

Thus, the principal adaptation from unconditional categorical sequence generation to protein inverse folding is the introduction of explicit geometric conditioning. The predicted amino-acid distribution at residue $i$ depends not only on its current simplex coordinates but also on the target backbone and the current probabilistic sequence state across the protein.

As in the denoising-classifier formulation of Dirichlet flow matching [1], the primary training objective is categorical cross-entropy against the native amino-acid identity. For the set $\mathcal{D}$ of positions included in the prediction objective,

$$\mathcal{L}_{\mathrm{AA}} = -\mathbb{E}_{\mathbf{B},\mathbf{y},t,\mathbf{X}_t}\left[\frac{1}{|\mathcal{D}|}\sum_{i\in\mathcal{D}}\log\widehat{p}_{\theta,i}\left(y_i \mid \mathbf{X}_t, \mathcal{G}(\mathbf{B}), t\right)\right]. \tag{S8}$$

Inverse FoldDir additionally uses auxiliary physicochemical prediction objectives corresponding to electrostatic or polarity classes and side-chain geometric classes. The complete training objective is

$$\mathcal{L}_{\mathrm{total}} = \mathcal{L}_{\mathrm{AA}} + \lambda_{\mathrm{elec}}\mathcal{L}_{\mathrm{elec}} + \lambda_{\mathrm{geom}}\mathcal{L}_{\mathrm{geom}}, \tag{S9}$$

with $\lambda_{\mathrm{elec}} = 0.25$ and $\lambda_{\mathrm{geom}} = 0.25$. These auxiliary terms are used only during training. Sequence generation is driven by the amino-acid prediction distribution in Equation S7. The auxiliary class definitions are provided in Supplementary Section S6, and the complete optimization procedure is summarized in Algorithm 2.

#### S1.4 From amino-acid predictions to a structure-conditioned flow

Dirichlet flow matching associates the conditional probability path in Equation S4 with a conditional vector field that transports points through the simplex toward a specified categorical vertex. Following the derivation of Stark et al. [1], the conditional vector field for amino-acid class $a$ can be written as

$$\mathbf{u}_t\left(\mathbf{x}_i \mid \mathbf{e}_a\right) = C(x_{i,a}, t)\left(\mathbf{e}_a - \mathbf{x}_i\right), \tag{S10}$$

where $C(x_{i,a}, t)$ is the position- and time-dependent scalar function derived for the Dirichlet probability path by Stark et al. The closed-form expression involves the regularized incomplete beta function. We use the previously derived Dirichlet flow field directly rather than rederiving it here.

The structure-conditioned marginal vector field at residue $i$ is obtained by averaging the class-specific conditional flows according to the amino-acid probabilities predicted by Inverse FoldDir:

$$\mathbf{v}_{\theta,i}\left(\mathbf{X}_t, \mathcal{G}(\mathbf{B}), t\right) = \sum_{a=1}^{K} \widehat{p}_{\theta,i}\left(a \mid \mathbf{X}_t, \mathcal{G}(\mathbf{B}), t\right) \mathbf{u}_t\left(\mathbf{x}_{i,t} \mid \mathbf{e}_a\right). \tag{S11}$$

Equation S11 connects amino-acid prediction to iterative sequence generation. The neural network does not directly regress an unconstrained vector in the simplex. Instead, it predicts the probability of each possible denoised amino-acid identity, and these probabilities determine the weighted combination of the corresponding Dirichlet conditional flows [1]. Because the probabilities in Equation S7 depend on the entire protein graph and current sequence state, the resulting flow at each residue is indirectly coupled to all other residues represented by the graph.

#### S1.5 Numerical sequence generation

Generation begins by assigning each unconstrained residue an initial state sampled from the uniform Dirichlet distribution,

$$\mathbf{x}_{i,0} \sim \mathrm{Dir}(\mathbf{1}). \tag{S12}$$

The learned vector field is then numerically integrated forward along the Dirichlet probability path. Here, $t$ denotes the Dirichlet time parameter, not normalized integration progress. Training times were drawn using exponential sampling, with a maximum Dirichlet time of $T = 100$. For $N$ uniformly spaced integration updates, normalized progress $s_k = k/N$ is mapped to Dirichlet time by

$$t_k = T s_k, \qquad \Delta t_k = T/N, \qquad k = 0, \ldots, N-1. \tag{S13}$$

Both model conditioning and the Dirichlet vector field use $t_k$ on this 0–100 scale. The integration increment is likewise expressed in Dirichlet time, so the terminal state is reached at $t_N = T$.

During sampling only, the amino-acid logits $z_{\theta,i,a}$ are divided by a temperature $\tau$ before softmax. We used $\tau = 1$ in the experiments reported here:

$$\widehat{p}^{(\tau)}_{\theta,i}(a \mid \mathbf{X}_t, \mathcal{G}(\mathbf{B}), t) = \frac{\exp(z_{\theta,i,a}/\tau)}{\sum_{b=1}^{K} \exp(z_{\theta,i,b}/\tau)}. \tag{S14}$$

These probabilities replace $\widehat{p}_{\theta,i}$ in Equation S11 to define the sampling vector field $\mathbf{v}^{(\tau)}_{\theta,i}$. At the temperature used here, $\tau = 1$, the logits are unchanged and $\mathbf{v}^{(1)}_{\theta,i} = \mathbf{v}_{\theta,i}$. Temperature scaling is not applied to the training loss, the Dirichlet corruption distribution, or the final argmax decoding. For integration step $k$,

$$\mathbf{x}_i^{(k+1)} = \Pi_\Delta \left[ \mathbf{x}_i^{(k)} + \Delta t_k \, \mathbf{v}_{\theta,i}^{(\tau)} \left( \mathbf{X}^{(k)}, \mathcal{G}(\mathbf{B}), t_k \right) \right], \tag{S15}$$

where $\Pi_\Delta$ denotes projection onto the probability simplex. All designable residue positions are updated during each integration step. Inverse FoldDir uses 20 denoising steps for the experiments reported here.

After the final integration step $N$, the continuous state is converted to a discrete protein sequence by selecting the highest-probability amino acid at each residue,

$$\widehat{y}_i = \arg\max_{a \in \{1,\ldots,K\}} x_{i,a}^{(N)}. \tag{S16}$$

Because every nonfixed residue remains represented by a complete probability distribution until the end of integration, its most probable identity may change between intermediate steps.

#### S1.6 Fixed residues and soft amino-acid priors

The simplex representation also permits sequence constraints to be incorporated directly into the generative state. For a set $\mathcal{F}$ of fixed residues with specified identities $c_i$, Inverse FoldDir sets

$$\mathbf{x}_i^{(k)} = \mathbf{e}_{c_i}, \qquad i \in \mathcal{F}, \tag{S17}$$

throughout generation. These positions therefore provide sequence context to the model but are not altered by the flow.

Soft residue priors use a different mechanism. For a position $i$ with user-defined prior distribution

$$\boldsymbol{\pi}_i \in \Delta^{K-1}, \tag{S18}$$

the initial state is set to

$$\mathbf{x}_i^{(0)} = \boldsymbol{\pi}_i \tag{S19}$$

rather than sampled from the uniform Dirichlet distribution. Unlike a fixed residue, this position remains designable during subsequent flow integration. A soft prior therefore specifies an initial preference over possible amino acids without requiring the final sequence to contain any particular residue.

Together, these formulations place unconstrained generation, fixed-residue inpainting, and soft residue-prior conditioning within the same structure-conditioned Dirichlet flow representation.

### S2 Nanobody design constraints and naive baselines

Complete sequences, constraint tiers, fixed-position annotations, sequence identity and similarity values, predicted structural-recovery metrics, and other design-level metadata for all tested nanobody constructs are provided in Supplementary Data 2.

#### S2.1 Tiers of fixed positions

The tiered constraints were selected prospectively as conservative safeguards for structural integrity or binding, before evaluating whether these residues would be recovered without explicit fixation. C23 and C97 were subsequently recovered in every generated Tier 0 (unconstrained) sequence. The anti-GFP nanobody designs used the following tiers of fixed positions:

**Tier 0:** None.

**Tier 1:** C23, C97, R36.

**Tier 2:** C23, C97, R36, E104, F103.

**Tier 3:** C23, C97, R36, E104, F103, N100, G102.

**Tier 4:** C23, C97, R36, E104, F103, N100, G102, S34, Y38.

**Tier 5:** C23, C97, R36, E104, F103, N100, G102, S34, Y38, W48, S60.

Blue text indicates residues newly fixed relative to the preceding tier.

#### S2.2 Naive baseline sequence generation

Our naive-baseline sequence-generation method simulated a scenario without generative AI guidance. Because Inverse FoldDir achieved an average native sequence recovery of approximately 55% for the nanobody, we generated naive-baseline sequences with the same approximate sequence identity by preserving 58 of the 105 positions. Eleven of these unchanged positions were the Tier-5 residues hypothesized to contribute to structural integrity or binding. Among the remaining

94 positions, 47 were randomly selected to remain unchanged and 47 were selected for mutation. We varied the conservativeness of these mutations as follows:

**Version 1: 55% sequence identity and 75% sequence similarity.**

- 55% identical, unmutated
- 11 positions fixed
- 47 additional positions randomly selected to remain unchanged among the 94 positions not fixed by Tier 5
- Remaining 45% mutated (47 positions out of 105)
- 20% conservative mutations with the highest BLOSUM alternative (21 positions out of the 47 positions randomly selected)
- 25% nonconservative mutations with negative BLOSUM scores (the remaining 26 positions)

**Version 2: 55% sequence identity and 100% sequence similarity.**

- 55% identical, unmutated
- 11 positions fixed
- 47 additional positions randomly selected to remain unchanged among the 94 positions not fixed by Tier 5
- Remaining 45% mutated (47 positions out of 105) to the alternative with the highest BLOSUM score

We generated 100 sequences for each version, predicted their structures, computed structural-recovery metrics, and selected representative candidates for experimental comparison.

### S3 Graph construction

#### S3.1 Residue-level node features

Table S1: Residue-level node features.

| Feature | GVP channel | Dim. | Description |
|---|---|---|---|
| Current amino-acid distribution | Scalar | 21 | Probability distribution over the 20 canonical amino acids and the unknown or masked class at denoising time $t$. |
| Backbone dihedral angles | Scalar | 6 | Sine and cosine encodings of $\phi$, $\psi$, and $\omega$, calculated from backbone N, C$\alpha$, and C coordinates. |
| Normalized structural uncertainty | Scalar | 1 | Within-protein normalized B-factor for experimentally determined structures or pLDDT for AlphaFold-predicted structures. Oriented so that higher values indicate more reliable geometry. Degenerate or unavailable values (e.g., NMR) were assigned the neutral value 0.5. |
| Structural source indicator | Scalar | 2 | Source encoding for experimentally determined and AlphaFold-predicted structures. |
| Geometry-missing indicator | Scalar | 1 | Binary feature identifying residues with missing or invalid backbone geometry. |
| Log packing density | Scalar | 1 | $\log(1+x)$, where $x$ is the Gaussian-weighted local packing score calculated from neighboring virtual C$\beta$ positions. |
| Burial percentile | Scalar | 1 | Within-structure percentile rank of the local packing score, providing a relative measure of residue burial. |
| Forward backbone direction | Vector | $1 \times 3$ | Normalized C$\alpha$-to-C$\alpha$ direction from residue $i$ to residue $i+1$. |
| Backward backbone direction | Vector | $1 \times 3$ | Normalized C$\alpha$-to-C$\alpha$ direction from residue $i$ to residue $i-1$. |
| Virtual C$\beta$ direction | Vector | $1 \times 3$ | Approximate side-chain direction inferred from backbone geometry using a virtual C$\beta$ atom. |

#### S3.2 Edge features

Table S2: Residue-graph edge features.

| Feature | GVP channel | Dim. | Description |
|---|---|---|---|
| Inter-residue distance encoding | Scalar | 16 | Radial basis expansion of the C$\alpha$-C$\alpha$ distance between connected residues. |
| Sequence-separation encoding | Scalar | 16 | Positional encoding of the signed residue-index difference $i - j$. |
| Relative direction | Vector | $1 \times 3$ | Unit vector from the source-residue C$\alpha$ coordinate to the target-residue C$\alpha$ coordinate. |

#### S3.3 Feature calculations

Table S3: Definitions of packing-density and burial features.

| Quantity | Definition | Interpretation |
|---|---|---|
| Raw packing score | $p_i = \sum_{\substack{j:\,\lvert i-j\rvert>2 \\ d_{ij}<14\,\text{Å}}} \exp\left[-\left(\frac{d_{ij}}{8.0\,\text{Å}}\right)^2\right]$ | Gaussian-weighted density of spatially neighboring residues around residue $i$. |
| Log packing density | $\log(1 + p_i)$ | Absolute local packing density with compressed dynamic range. |
| Burial percentile | Percentile rank of $p_i$ within the same structure | Relative burial of a residue compared with other residues in the same protein. |

#### S3.4 Handling of unresolved coordinates

Residues with missing or nonfinite backbone coordinates were retained as sequence positions but masked from geometric interactions and training losses. Missing-coordinate residues did not participate in distance-based graph connectivity or contribute geometric edge features, whereas sequence-adjacent edges were retained to preserve chain connectivity. An explicit missing-geometry indicator was provided as a node feature.

### S4 Training dataset construction

#### S4.1 CATH 4.2 splits

We used the original CATH 4.2 chain-level splits released by Ingraham et al. [3] without re-splitting the data. The splits contained 18,024 training chains, 608 validation chains, and 1,120 test chains. Only the 18,024-chain training split was used for model fitting. The validation and test splits remained held out for model selection and final evaluation, respectively.

#### S4.2 AlphaFoldDB cluster filtering and redundancy reduction

The predicted-structure training set was constructed from AlphaFoldDB [4] using the structural-cluster release of Barrio-Hernandez et al. [5]. We began with 2,302,907 Foldseek-based clusters containing at least two members. Structural redundancy was controlled using these published clusters rather than by treating every AlphaFoldDB structure as an independent training example. Clusters were retained only when both the representative-structure pLDDT and the mean pLDDT across cluster members exceeded 70.

##### S4.3 Structural filtering of AlphaFoldDB training clusters

Using available SIFTS mappings [6], we linked held-out CATH 4.2 validation and test chains to AlphaFoldDB cluster representatives. We used Foldseek [7] to compare these representatives with candidate training-cluster representatives and excluded entire training clusters when a match had $E < 0.1$.

After cluster filtering, we retained individual AlphaFoldDB structures with mean pLDDT $\geq 70$ and sequence length $\leq 700$ residues, yielding 2,824,736 predicted structures. During training, AlphaFoldDB-predicted and experimentally determined CATH structures were sampled at a 1:1 ratio.

#### S5 Hyperparameters

##### S5.1 Training

**Epochs:** 3000

**Training time:** 7 days

**Training hardware:** One NVIDIA L40S GPU with 48 GB VRAM

**Learning rate:** $5 \times 10^{-4}$

**Batch size:** 20

**Optimizer:** AdamW

**Learning rate scheduler:** ReduceLROnPlateau

**Dropout:** 0

**Weight decay:** 0.01

**GVP layers:** 5

**Message passing:** 4

**Prediction head layers:** 4

**Hidden scalar dimension:** 288

**Hidden vector dimension:** 32

**Prediction head hidden dimension:** 128

**Time embedding dimension:** 64

**Time embedding integration:** FiLM

**Target-class Dirichlet concentration:** $1 + t$, ranging from 1 to 101; the other classes have concentration 1 (Equation S4).

**Time sampling strategy:** Exponential sampling with maximum Dirichlet time $T = 100$.

**Maximum C$\alpha$ distance for edge formation:** 12 Å

**Radial basis function dimension for 3D distance:** 16

**Radial basis function dimension for sequence distance:** 16

**Ratio of AlphaFold2-predicted structures to experimentally determined structures in the training set:** 1

**Standard deviation of the Gaussian noise on atom coordinates:** 0.25 Å

**Lambda electrostatic loss:** 0.25

**Lambda geometry/topology loss:** 0.25

**Ratio of batches with mixed training mode:** 50%

Here, an epoch was defined relative to the experimentally determined CATH 4.2 training split, not the complete combined training corpus. One epoch corresponded to one traversal of the 18,024 CATH training chains, with an equal number of AlphaFoldDB structures sampled during the same interval to maintain the 1:1 source ratio. Thus, an epoch contained approximately 36,048 sampled structures, corresponding to 1,802 full minibatches and a final partial minibatch at a batch size of 20. It did not represent a pass through all 2,824,736 AlphaFoldDB structures.

#### S5.2 Training procedure

The complete training procedure is summarized in Algorithm 2.

---

**Algorithm 2** Training algorithm for Inverse FoldDir

---

**Require:** training proteins with backbone structures and native sequences; model parameters $\theta$

**for all** training batches **do**
- Sample a protein backbone $B$ and its native sequence $Y$
- $\widetilde{B} \leftarrow$ PERTURBCOORDINATES$(B)$
- $G \leftarrow$ CONSTRUCTRESIDUEGRAPH$(\widetilde{B})$
- Sample a denoising time $t$ using the exponential time-sampling strategy
- $X_t \sim$ CONDITIONALDIRICHLETPATH$(Y, t)$
- Optionally construct mixed-generation or inpainting conditions for $X_t$
- $(\widehat{P}, \widehat{Q}_{\text{elec}}, \widehat{Q}_{\text{geom}}) \leftarrow$ INVERSE FOLDDIR$()\theta G, X_t, t$
- $\mathcal{L}_{\text{AA}} \leftarrow$ AMINOACIDLOSS$(\widehat{P}, Y)$
- $\mathcal{L}_{\text{elec}} \leftarrow$ ELECTROSTATICAUXILIARYLOSS$(\widehat{Q}_{\text{elec}}, Y)$
- $\mathcal{L}_{\text{geom}} \leftarrow$ GEOMETRICAUXILIARYLOSS$(\widehat{Q}_{\text{geom}}, Y)$
- $\mathcal{L} \leftarrow \mathcal{L}_{\text{AA}} + \lambda_{\text{elec}}\mathcal{L}_{\text{elec}} + \lambda_{\text{geom}}\mathcal{L}_{\text{geom}}$
- $\theta \leftarrow$ OPTIMIZERSTEP$(\theta, \nabla_\theta \mathcal{L})$

**end for**

---

#### S5.3 Inference

**Steps:** 20

**Maximum Dirichlet time:** $T = 100$; normalized integration progress is mapped to Dirichlet time by $t = Ts$.

**Sampling temperature:** $\tau = 1$, leaving amino-acid logits unchanged before softmax at each generation update (Equation S14); not used during training.

**Standard deviation of the Gaussian noise on atom coordinates:** 0 Å

### S6 Auxiliary prediction heads

Table S4: Auxiliary prediction heads used during training.

| **Auxiliary head** | **Property category** | **Classes** | **Amino-acid grouping** | **Loss weight** |
|---|---|---|---|---|
| Electrostatic and polarity class | Physicochemical | 5 | Acidic: D, E; basic: K, R; neutral polar: N, Q, S, T; histidine: H; other: remaining residues | 0.25 |
| Geometric topology class | Side-chain shape and steric packing | 5 | Aromatic: F, W, Y; C$\beta$-branched: I, V; extended: L, M; small: A, G; other: remaining residues | 0.25 |

### S7 Experimental methods

#### S7.1 Experimental sample mapping

The complete computational design inventory is provided in Supplementary Data 2.

#### S7.2 Cell-free protein expression

Crude extracts were prepared from *Escherichia coli* BL21(DE3*) following previously reported cell-free extract preparation protocols [8, 9, 10]. All anti-GFP nanobodies and negative baselines were expressed in 7.5-$\mu$L cell-free reactions in a 96-well plate. sfGFP-6$\times$His was expressed in 15-$\mu$L cell-free reactions in 2-mL microcentrifuge tubes. Both sfGFP and the nanobodies were incubated overnight at room temperature using the PANOx-SP formulation [11].

To support the production of folded nanobodies containing disulfide bonds, an oxidizing reaction environment was established based on prior cell-free production of disulfide-bonded proteins [12]. Cell extracts were pretreated with 25 $\mu$M iodoacetamide (IAM) at room temperature for 30 min before use. An additional 4 mM oxidized glutathione (GSSG), 1 mM reduced glutathione (GSH), and 5 $\mu$M purified DsbC were added to the reaction mixture to further facilitate nanobody formation.

#### S7.3 AlphaLISA binding assay

Binding mixtures containing the nanobodies and sfGFP were diluted in a buffer consisting of 50 mM HEPES at pH 7.4, 150 mM NaCl, 1 mg/mL BSA, and 0.015% v/v Triton X-100. AlphaLISA experiments [13] used anti-FLAG donor beads and anti-6×His acceptor beads.

Following dilution, an Echo 525 acoustic liquid handler was used to transfer 0.5 $\mu$L of cell-free-expressed sfGFP-6×His, 0.5 $\mu$L of cell-free-expressed anti-GFP nanobody, 0.5 $\mu$L of blank buffer, and 0.25 $\mu$L of anti-FLAG donor beads diluted in buffer from a 384-well polypropylene 2.0 Plus Source microplate to an AlphaPlate 1536-well destination microplate (Revvity) using the 384PP_Plus_GPSA fluid type. The plate was sealed and equilibrated for 1 h at room temperature. Following incubation, 0.125 $\mu$L of anti-6×His acceptor beads diluted in buffer was transferred to each reaction. The reactions were equilibrated for an additional hour at room temperature in the dark. For analysis, the reactions were incubated for 10 min in a Biotek Synergy Neo2 plate reader at room temperature, and the chemiluminescent signal was measured using the AlphaLISA filter with an excitation time of 100 ms, an integration time of 300 ms, and a settle time of 20 ms. An appropriate dilution factor was first determined using a dilution series of the wild-type nanobody and sfGFP-6×His in the AlphaLISA assay. The remaining nanobodies were then diluted accordingly and assayed. Protein concentrations were normalized based on the soluble yield of each design, as determined using a $^{14}$C-leucine incorporation assay [14].

#### S7.4 AlphaLISA data analysis

AlphaLISA signal was normalized to the mean of five non-binding controls: four naive-baseline sequences and the no-template control. Control variability pooled across the analyzed measurements was used to define a background threshold of 1.74-fold over the normalized background, corresponding to three standard deviations above the control signal. Designs exceeding this threshold in all four measurements were classified as confirmed binders. Designs exceeding the threshold in three of four measurements were classified as candidate binders.

Designs #27 and #29 satisfied the confirmed-binder criterion, with geometric-mean signals of 2.96- and 2.94-fold over background, respectively. Designs #13, #31, and #34 satisfied the candidate-binder criterion. The wild-type anti-GFP nanobody produced a 27.4-fold signal over background, whereas the five non-binding controls remained near background. The identities of the two confirmed binders were unchanged under alternative control definitions and threshold calculations.

##### S7.5 Boltz-2 cofolding analysis

We evaluated the redesigned nanobody-sfGFP pairs using Boltz-2 [15] as an orthogonal computational control. Across the 35 Inverse FoldDir designs, whole-complex confidence did not correlate with AlphaLISA signal either with an MSA (Spearman $\rho = 0.06$, $P = 0.74$) or in single-sequence mode ($\rho = -0.03$, $P = 0.86$; Figure S4). Boltz-2 ipTM ($\rho = 0.17$, $P = 0.34$) and complex pLDDT ($\rho = 0.00$, $P = 1.00$) likewise showed no significant association with experimental binding. ESMFold self-consistency TM-score, which was used during candidate selection, was also not significantly associated with binding signal ($\rho = -0.28$, $P = 0.10$; Figure S5). These results indicate that the evaluated computational structure-confidence metrics did not reliably rank experimentally retained binding within this design set.

#### S8 Ablation experiments

We examined which components of Inverse FoldDir contributed most strongly to structural recovery by training ablated models under the same self-consistency benchmark while removing individual training or architectural components.

Removing the structurally filtered AlphaFold-predicted training structures produced the largest performance decrease, reducing TM-score and increasing C$\alpha$ RMSD relative to the full model (Table 3). This result indicates that increasing the structural diversity of the training set contributed substantially to final performance.

Removing iterative denoising updates also significantly reduced structural recovery, whereas removing the auxiliary charge and steric prediction heads, mixed-mode training, entropy features, or burial descriptors produced only modest changes (Table 3). Together, these ablations suggest that training-data breadth and iterative refinement contributed more strongly than the individual auxiliary components evaluated here.

### S9 Baseline methods and evaluation protocols

We evaluated five published inverse-folding models (ProteinMPNN [16], MapDiff [17], FAMPNN [18], ESM-IF1 [19], and PiFold [20]) for structure recovery on a held-out set of 1,120 single-chain PDB structures, with applicable baselines also evaluated for zero-shot fitness prediction on single-amino-acid substitution variants from a curated subset of 35 ProteinGym deep-mutational-scanning assays [21]. Each baseline was run in a separate environment using the software versions required by its original repository. We used the publicly released pretrained checkpoints without retraining. Their original training datasets differed. ESM-IF1, for example, was trained using experimental structures and approximately 12 million AlphaFold2-predicted structures [19]. The benchmark therefore compares released models under the specified inference settings and a shared refolding and structure-comparison pipeline, rather than architectures trained on an identical dataset.

#### S9.1 Structure recovery

##### S9.1.1 Test set

The test set contains 1,120 single-chain PDB structures reduced to backbone atoms. Preprocessing retained only the primary alternate conformation, removed hetero-atoms and solvent molecules, used the first model in files containing multiple models, and incorporated insertion codes into integer residue numbering. Chain lengths range from 40 to 497 residues, with a median length of 136.

##### S9.1.2 Sampling

For each model and PDB structure, we generated three sequence designs. A sampling temperature of 0.3 was used for ProteinMPNN, FAMPNN, ESM-IF1, and PiFold. MapDiff was run using its default discrete-diffusion sampling procedure.

ProteinMPNN and ESM-IF1 were sampled through three independent autoregressive decoding runs. FAMPNN was run independently three times using its autoregressive sequence-generation and side-chain denoising procedure. PiFold produces one amino-acid distribution per position in a single forward pass. Because the original implementation returns a deterministic argmax sequence, we sampled three designs from the per-position logits using temperature-scaled multinomial sampling.

Reported means were calculated across all 1,120 structures. Standard deviations were calculated across the three design replicates after averaging each replicate over the complete test set. For each method and protein, each structural-recovery metric was averaged across the three generation replicates. The full-test-set $t$-tests used these 1,120 per-protein means per method, rather than the three test-set replicate means or the 3,360 individual designs.

##### S9.1.3 Folding

Each designed sequence was folded using ESMFold v1 [22]. The language-model trunk was run in half precision, while the structure module was run in single precision.

##### S9.1.4 Evaluation

Each predicted structure was compared with its corresponding reference structure using TM-align [23]. The primary metric was the minimum of the two TM-scores obtained by normalizing with respect to the lengths of chain 1 and chain 2.

The structure-comparison pipeline retained only C$\alpha$ atoms with a blank or "A" alternate-location indicator. Without this filter, alternate conformations in the reference structures could be counted more than once when determining chain length, affecting the length-normalized TM-score.

##### S9.1.5 Model configurations

Table S5: Baseline model configurations and sampling procedures.

| **Model** | **Weights** | **Temperature / seed** | **Sampling procedure** |
|---|---|---|---|
| ProteinMPNN (Dauparas et al., 2022) | Vanilla 48-neighbor checkpoint (0.20 Å C$\alpha$ noise) | 0.3 / 13 | Random-order autoregressive decoding |
| MapDiff (Bai et al., 2025) | Released checkpoint | / 34 | DDIM, 100 steps; 50-sample Monte Carlo dropout ensemble |

| Model | Weights | Temperature / seed | Sampling procedure |
|---|---|---|---|
| FAMPNN (Widatalla et al., 2025) | CATH-pretrained checkpoint (0.3 Å noise) | 0.30 / 37 | Autoregressive sequence sampling with side-chain denoising; 100 sequence and 50 side-chain steps |
| ESM-IF1 (Hsu et al., 2022) | Released 142M GVP-Transformer checkpoint (UR50 and AlphaFold structures) | 0.3 / 13 | Left-to-right autoregressive decoding |
| PiFold (Gao, Tan, and Li, 2023) | Released checkpoint | 0.3 / 13 | One-shot decoding with temperature-scaled multinomial sampling |

Each model produced 3,360 designs, corresponding to three designs for each of the 1,120 test structures.

#### S9.2 Zero-shot fitness prediction on ProteinGym

##### S9.2.1 Scoring setup

We evaluated the applicable baselines on ProteinGym [21] using an inpainting-style scoring procedure. Given the wild-type backbone and the wild-type residues at all unmutated positions, each model assigns a conditional log-likelihood ratio between the mutant and wild-type amino acids at the mutated positions.

This differs from the full-sequence procedure used in the original ProteinGym baseline scripts, in which the complete mutant sequence is teacher-forced and scored using its mean per-residue log-likelihood. For ProteinMPNN and ESM-IF1, we report results using both scoring procedures.

##### S9.2.2 Assay selection

We began with the 217 substitution assays in the ProteinGym reference table and retained assays in the ProteinGym “Medium” MSA-depth category.

From this category, we selected 35 assays covering the three broad ProteinGym selection types: 20 Activity assays, 13 Stability assays, and 2 Binding assays. Of these source assays, 24 contain only single substitutions and 11 also contain multi-mutant variants. Only single-substitution variants were retained for benchmarking in all 35 assays. Multi-mutant variants were excluded from every scoring procedure.

Protein lengths range from 37 to 1,159 residues, with a median length of 281. The source organisms include human in 24 assays, mouse in 5, yeast in 2, Arabidopsis in 2, one reconstructed ancestral protein, and one Lipomyces starkeyi enzyme.

##### S9.2.3 Structural input

For each assay, the ProteinGym-provided AlphaFold2 structure [24] was used as the structural input. Wild-type sequences were read directly from the corresponding structures to maintain alignment between sequence and structure indices.

##### S9.2.4 Inpainting-mode scoring

**ProteinMPNN** For each mutated position, ProteinMPNN was provided with the backbone, the wild-type sequence, and a design mask marking only the queried position as designable. The queried position was decoded last, so its predicted amino-acid distribution was conditioned on the backbone and on all other positions being fixed to their wild-type residues.

The mutation score was the log-likelihood ratio between the mutant and wild-type amino acids.

**ESM-IF1** ESM-IF1 is a left-to-right autoregressive model. Its amino-acid distribution at position i has the form

$$p(x_i \mid x_{<i}, \text{backbone}),$$

and is therefore conditioned on the backbone and on residues preceding position i.

A forward pass was performed using the wild-type sequence, and the distribution at each mutated position was used to calculate the log-likelihood ratio between the mutant and wild-type amino acids.

**FAMPNN** For all evaluated assays, we used FAMPNN's exhaustive single-mutant scorer. This procedure produces a position-by-amino-acid score matrix normalized relative to the wild-type amino acid at each position. The score corresponding to each mutation was selected from this matrix.

This procedure used the CATH-pretrained checkpoint recommended for mutation scoring.

**Excluded baselines** MapDiff and PiFold were not evaluated on the ProteinGym task.

The released MapDiff implementation does not provide a per-position mutation-scoring procedure. Its sequence predictions are obtained through the complete discrete-diffusion sampling process and cannot be directly converted into the conditional mutation scores used in this evaluation.

PiFold produces per-position amino-acid distributions conditioned on the backbone, but not on the wild-type residues at the remaining positions. It therefore does not represent the conditional distribution required for the inpainting evaluation.

###### S9.2.5 Full-sequence scoring

**ProteinMPNN** The wild-type backbone and complete mutant sequence were passed to the model in a single forward pass. The variant score was the mean negative log-likelihood across the sequence. Only single-substitution variants were evaluated.

**ESM-IF1** The complete mutant sequence was teacher-forced conditional on the wild-type backbone. The variant score was the average per-residue log-likelihood. Only single-substitution variants were evaluated.

FAMPNN was evaluated only using the mutation-scoring procedures described above under *Inpainting-mode scoring*.

###### S9.2.6 Metric

For each assay and model-score threshold, we calculated positive predictive value (PPV) as the fraction of score-selected variants classified as experimental positives. Experimental positives were defined using the assay-specific binary labels provided in ProteinGym's preprocessed tables, which encode ProteinGym's binarization of the experimental DMS values. We therefore did not apply a single universal DMS-score threshold across assays. Only variants with finite model scores were included.

### S10 Supplementary figures

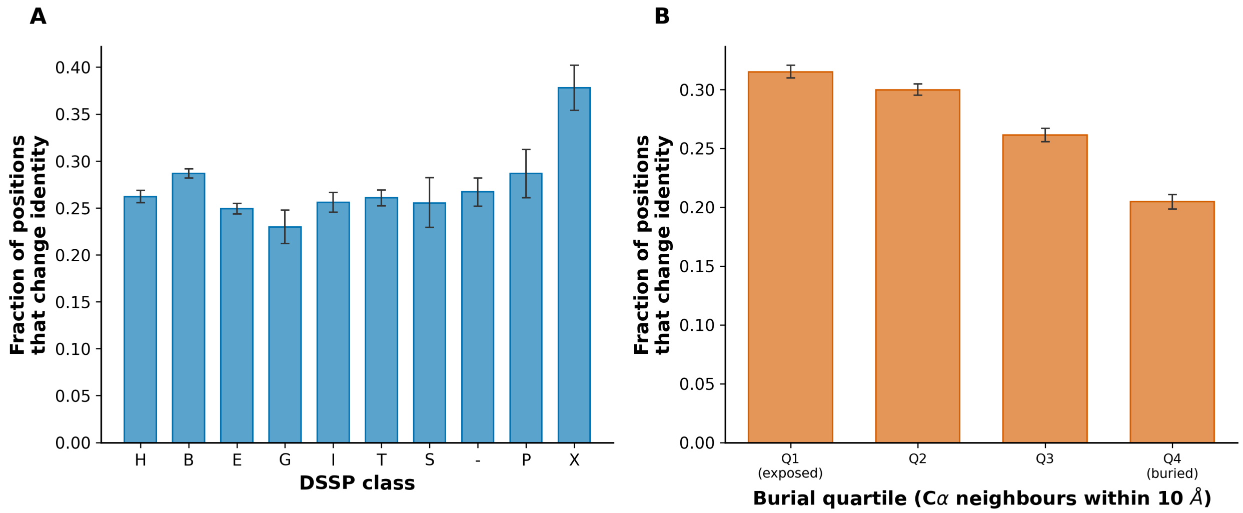

Figure S1: Residue identity changes during denoising across structural environments. (A) Fraction of positions that change identity by DSSP secondary-structure class [25]. (B) Fraction of positions that change identity by residue-burial quartile, defined using the number of C$\alpha$ neighbors within 10 Å. Error bars show variability across proteins.

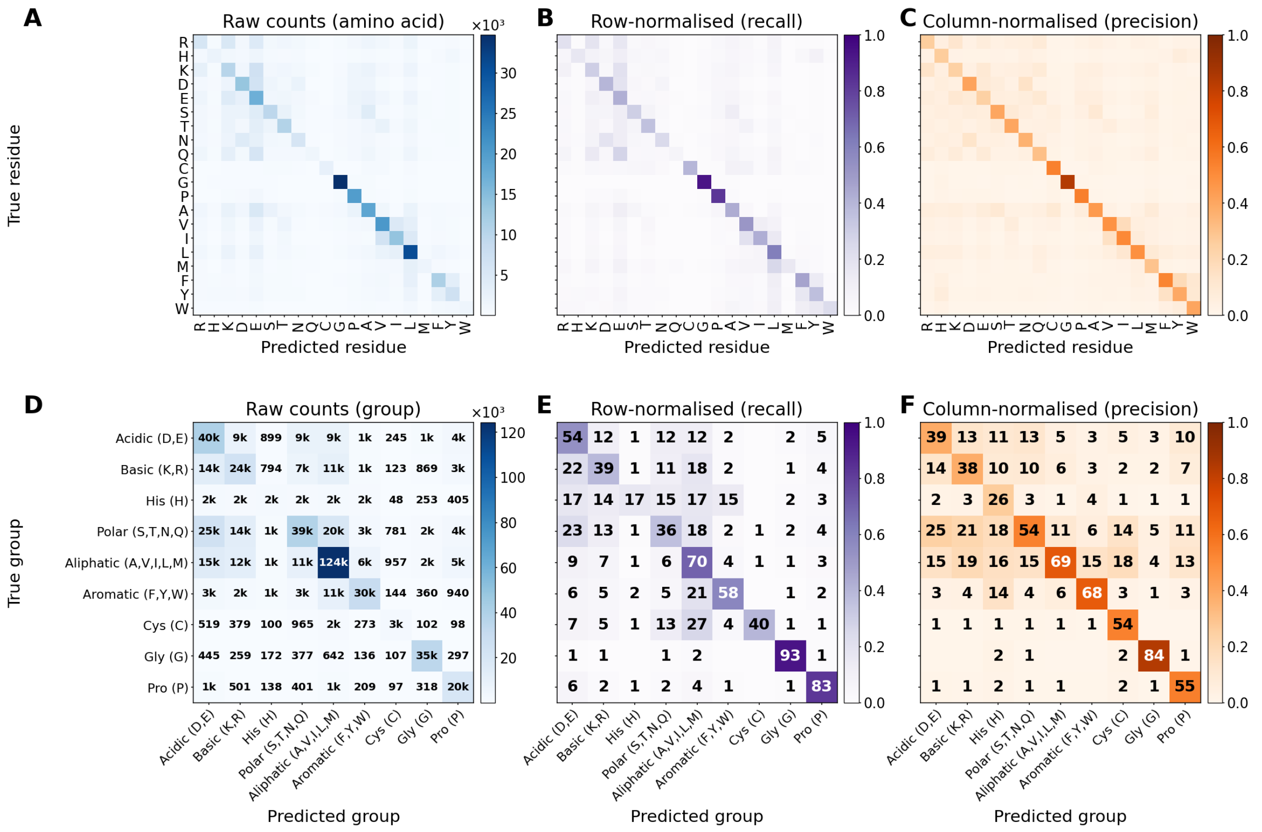

Figure S2: Final-sequence amino-acid recovery patterns on the CATH 4.2 test set. Confusion matrices compare native residues or residue groups (rows) with residues predicted in the final generated sequences (columns), aggregated across the three generation replicates. (A-C) Amino-acid-level raw counts, row-normalized recall, and column-normalized precision, respectively. (D-F) Corresponding matrices after grouping amino acids into broad physicochemical or conformational classes: acidic (D, E), basic (K, R), histidine (H), polar (S, T, N, Q), aliphatic (A, V, I, L, M), aromatic (F, Y, W), cysteine (C), glycine (G), and proline (P). Values in the grouped raw-count matrix are shown as counts, abbreviated with “k” for thousands. Values in the normalized grouped matrices are percentages. Row normalization shows the distribution of predictions for each native residue or group, whereas column normalization shows the composition of each predicted residue or group. The grouped matrices show preferential recovery within hydrophobic classes and particularly high identity-level recovery for glycine and proline.

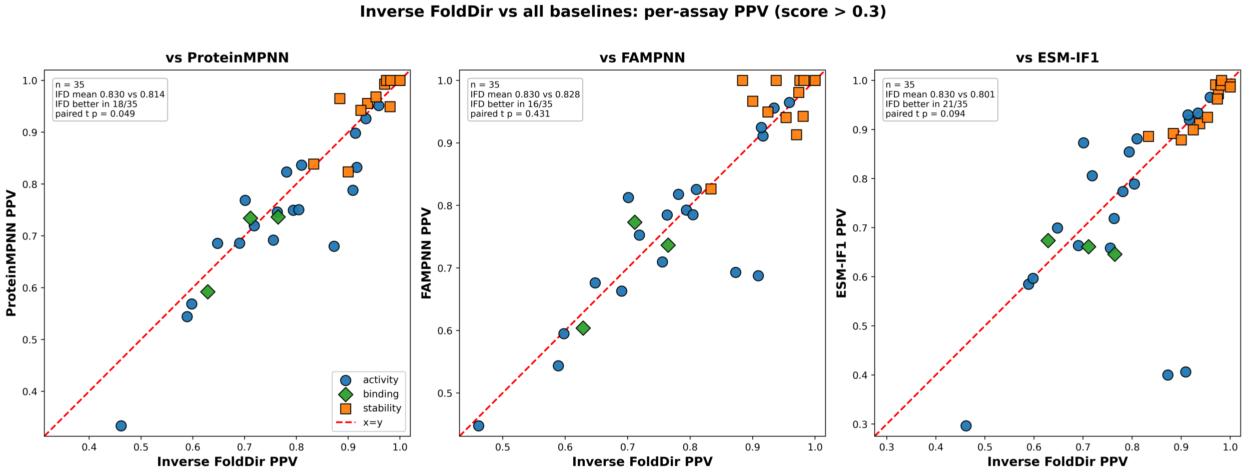

Figure S3: Per-assay ProteinGym [21] PPV comparisons between Inverse FoldDir and ProteinMPNN, FAMPNN, and ESM-IF1 at a score threshold greater than 0.3. Points are grouped by activity, binding, and stability assays. The dashed line indicates equal performance.

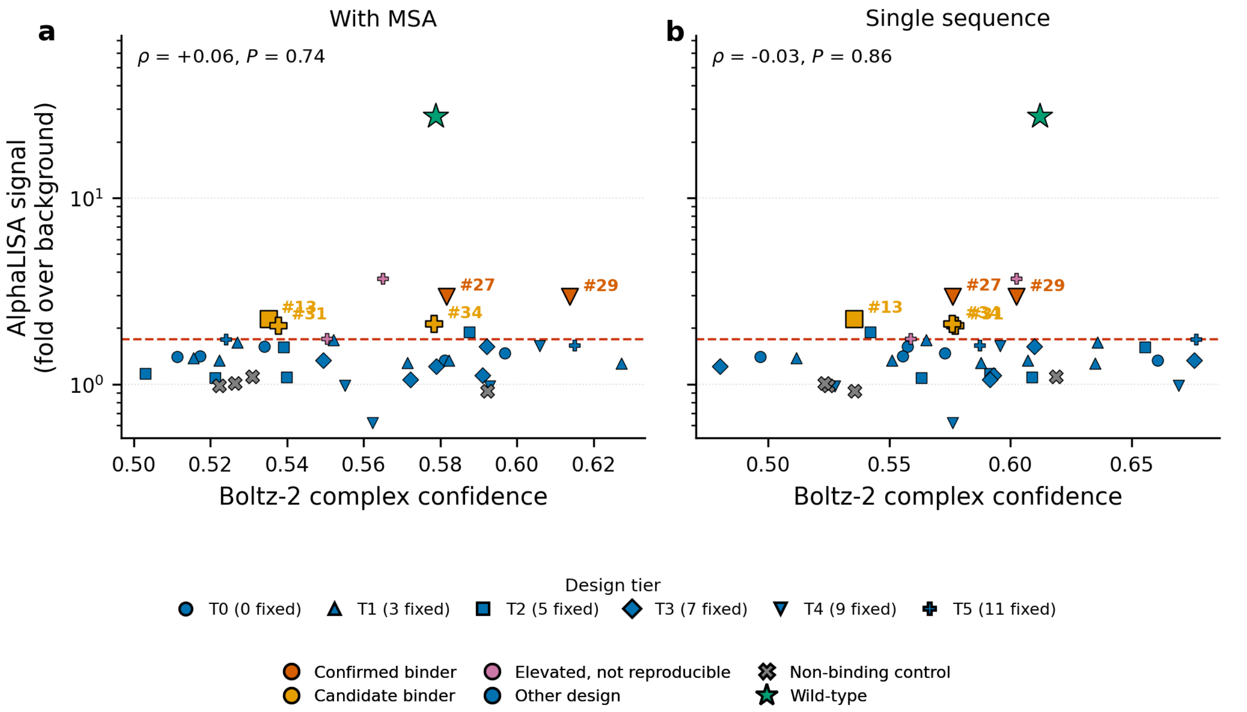

Figure S4: Boltz-2 cofolding confidence compared with experimental binding signal. Each design was cofolded with sfGFP using Boltz-2 v2.0.3 with a multiple-sequence alignment (A) and in single-sequence mode (B). The $x$-axis shows the Boltz-2 `confidence_score`, defined for a multichain complex as $0.8\times$ complex pLDDT plus $0.2\times$ ipTM, with both components on a 0-1 scale. This score combines average local structural confidence across the complex with interface confidence, rather than measuring interface confidence alone. The $y$-axis shows the replicate-matched AlphaLISA signal. Marker shape denotes the conservation tier (number of fixed residues), and color denotes the binding classification used in Figure 6. The wild-type nanobody (star) and non-binding controls (crosses) are shown as experimental anchors but were excluded from the correlations, which were computed across the 35 designs.

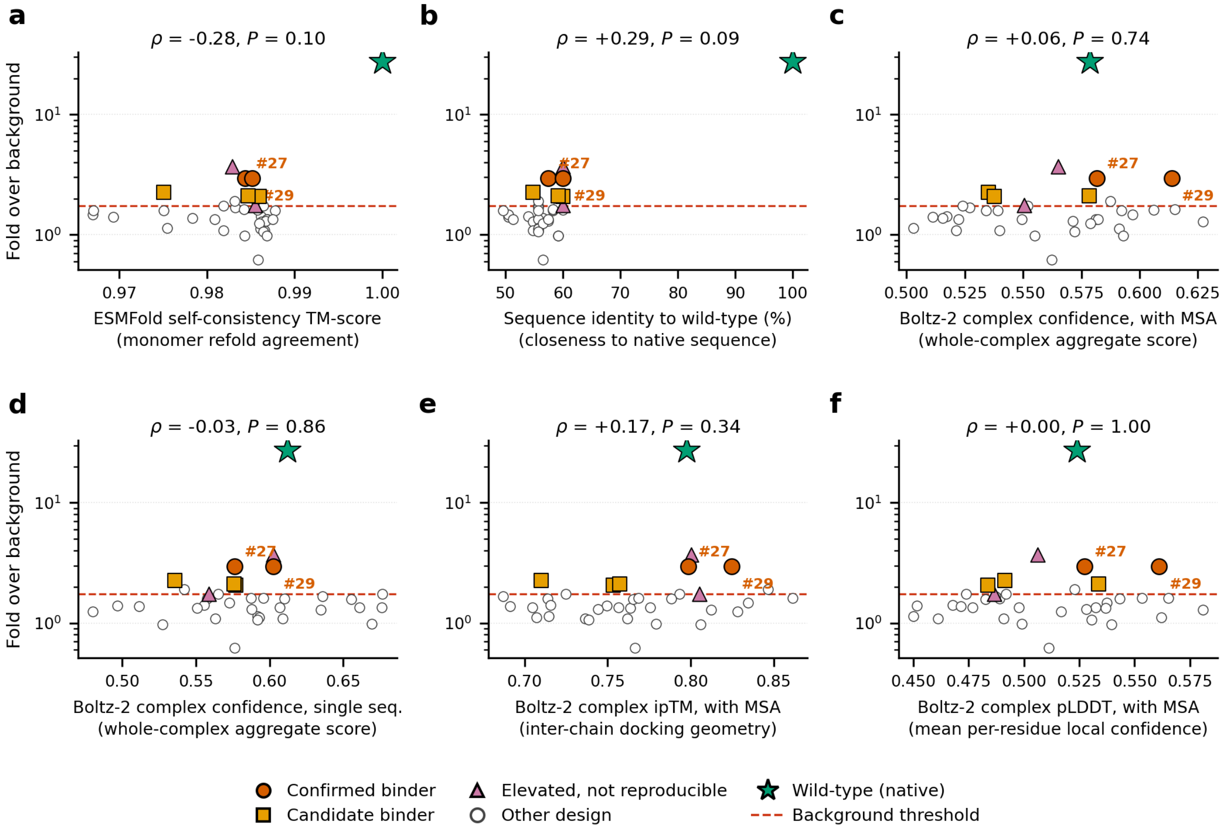

Figure S5: Computational predictors compared with experimental binding signal. Each panel compares one computational predictor with the replicate-matched AlphaLISA signal across the 35 designs. Spearman $\rho$ and $P$ values are shown above each panel and were computed using designs only. Construct classes are distinguished by both marker shape and color to preserve readability in grayscale: filled circles, confirmed binders; filled squares, candidate binders; filled triangles, elevated but not reproducible; open circles, other designs; and star, wild-type nanobody. The dashed line indicates the background threshold, and each $x$-axis sublabel states what the corresponding metric measures. The wild-type (native) sequence is shown in every panel as a reference but was excluded from the correlations.

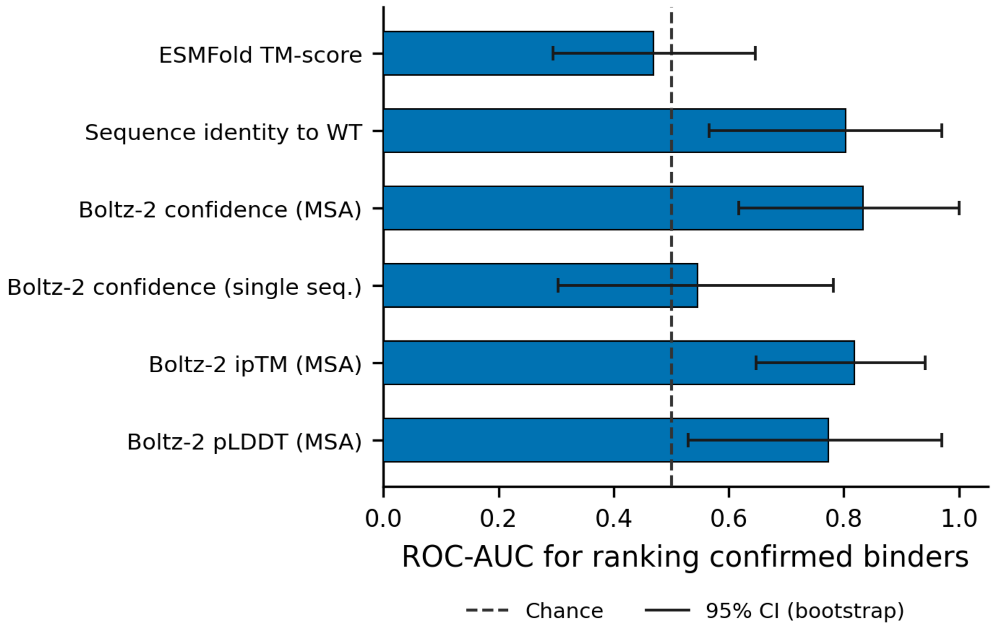

Figure S6: Discrimination of confirmed nanobody binders by computational predictors. Bars show the area under the receiver operating characteristic curve (ROC-AUC) for ranking the confirmed binders above the remaining designs. Error bars indicate 95% percentile-bootstrap confidence intervals from 10,000 resamples of the 35 designs. Resamples containing no confirmed binder were undefined and excluded. Every interval includes or nearly includes chance performance (0.5), and no predictor significantly enriched confirmed binders among its eight highest-ranked designs (hypergeometric $P \geq 0.41$ for all six predictors). With only two confirmed binders, the point estimates are not statistically separable and should not be used to rank predictors.

### References

[1] Hannes Stark, Bowen Jing, Chenyu Wang, Gabriele Corso, Bonnie Berger, Regina Barzilay, and Tommi Jaakkola. Dirichlet flow matching with applications to dna sequence design. *arXiv preprint arXiv:2402.05841*, 2024.

[2] Pavel Avdeyev, Chenlai Shi, Yuhao Tan, Kseniia Dudnyk, and Jian Zhou. Dirichlet diffusion score model for biological sequence generation. In *International Conference on Machine Learning*, pages 1276–1301. PMLR, 2023.

[3] John Ingraham, Vikas Garg, Regina Barzilay, and Tommi Jaakkola. Generative models for graph-based protein design. *Advances in neural information processing systems*, 32, 2019.

[4] Mihaly Varadi, Stephen Anyango, Mandar Deshpande, Sreenath Nair, Cindy Natassia, Galabina Yordanova, David Yuan, Oana Stroe, Gemma Wood, Agata Laydon, et al. AlphaFold Protein Structure Database: massively expanding the structural coverage of protein-sequence space with high-accuracy models. *Nucleic Acids Research*, 50(D1):D439–D444, 2022.

[5] Inigo Barrio-Hernandez, Jingi Yeo, Jürgen Jänes, Milot Mirdita, Cameron LM Gilchrist, Tanita Wein, Mihaly Varadi, Sameer Velankar, Pedro Beltrao, and Martin Steinegger. Clustering predicted structures at the scale of the known protein universe. *Nature*, 622(7983):637–645, 2023.

[6] Jose M Dana, Aleksandras Gutmanas, Nidhi Tyagi, Guoying Qi, Claire O'Donovan, Maria Martin, and Sameer Velankar. Sifts: updated structure integration with function, taxonomy and sequences resource allows 40-fold increase in coverage of structure-based annotations for proteins. *Nucleic acids research*, 47(D1):D482–D489, 2019.

[13] Edwin F Ullman, Hrair Kirakossian, Singh Singh, Z Ping Wu, Benjamin R Irvin, John S Pease, Arthur C Switchenko, Jennifer D Irvine, Alan Dafforn, and Carl N Skold. Luminescent oxygen

channeling immunoassay: measurement of particle binding kinetics by chemiluminescence. *Proceedings of the National Academy of Sciences*, 91(12):5426–5430, 1994.

[14] Rusty J Mans and G David Novelli. Measurement of the incorporation of radioactive amino acids into protein by a filter-paper disk method. *Archives of Biochemistry and Biophysics*, 94(1):48–53, 1961.

[15] Saro Passaro, Gabriele Corso, Jeremy Wohlwend, Mateo Reveiz, Stephan Thaler, Vignesh Ram Somnath, Noah Getz, Tally Portnoi, Julien Roy, Hannes Stark, et al. Boltz-2: Towards accurate and efficient binding affinity prediction. *BioRxiv*, 2025.

[16] Justas Dauparas, Ivan Anishchenko, Nathaniel Bennett, Hua Bai, Robert J. Ragotte, Lukas F. Milles, Basile I. M. Wicky, Alexis Courbet, Rob J. de Haas, Neville Bethel, et al. Robust deep learning–based protein sequence design using proteinmpnn. *Science*, 378(6615):49–56, 2022.

[17] Peizhen Bai, Filip Miljković, Xianyuan Liu, Leonardo De Maria, Rebecca Croasdale-Wood, Owen Rackham, and Haiping Lu. Mask-prior-guided denoising diffusion improves inverse protein folding. *Nature Machine Intelligence*, pages 1–13, 2025.

[18] Talal Widatalla, Richard W Shuai, Brian Hie, and Possu Huang. Sidechain conditioning and modeling for full-atom protein sequence design with fampnn. In *Forty-second International Conference on Machine Learning*, 2025.

[19] Chloe Hsu, Robert Verkuil, Jason Liu, Zeming Lin, Brian Hie, Tom Sercu, Adam Lerer, and Alexander Rives. Learning inverse folding from millions of predicted structures. In *International conference on machine learning*, pages 8946–8970. PMLR, 2022.

[20] Zhangyang Gao, Cheng Tan, Pablo Chacón, and Stan Z Li. Pifold: Toward effective and efficient protein inverse folding. *arXiv preprint arXiv:2209.12643*, 2022.
